# Massively parallel identification of single-cell immunophenotypes

**DOI:** 10.1101/2024.04.04.587924

**Authors:** Martin Cienciala, Laura Alvarez, Laura Berne, David Chena, Pavel Fikar, Monika Holubova, Hynek Kasl, Daniel Lysak, Mona Luo, Zuzana Novackova, Sheyla Ordonez, Zuzana Sramkova, Tomas Vlas, Daniel Georgiev

## Abstract

Translating insights from single-cell analysis into actionable indicators of health and disease requires large-scale confirmatory studies. We introduce biocytometry, a novel method utilizing engineered bioparticles for multiparametric immunophenotyping in suspension, enabling simultaneous measurement across thousands of assays with single-cell sensitivity and a wide dynamic range (1 to 1,000 target cells/sample). The technical validation of biocytometry revealed strong alignment with established technologies (mean bias = 0.25%, LoA = −1.83% to 2.33%) for low-sensitivity settings. Biocytometry excelled in high-sensitivity settings, consistently showcasing superior sensitivity and specificity (LoB = 0), irrespective of the sample type. By employing multiparametric target cell identification, we harnessed the homogeneous assay workflow to discern cell-specific apoptosis in mixed cell cultures. Potential applications include monitoring rare premalignant subpopulations in indications such as smoldering multiple myeloma (SMM), enhancing the detection of circulating tumor cells (CTCs), advancing pharmacokinetic assessments in chimeric antigen receptor (CAR) T-cell therapies, and improving the accuracy of minimal residual disease (MRD) evaluations. Additionally, the high throughput and cell-specific readout capabilities might provide substantial value in drug development, especially for the analysis of complex sample matrices, such as primary cell cultures and organoids.

## Introduction

Single-cell research sits at the intersection of our understanding of health and disease, offering unprecedented insights into cellular mechanisms and creating opportunities throughout the pharmaceutical value chain^1–3^. However, the swift translation of these discoveries into real-world applications is hampered by significant limitations of current technologies^4–7^. Chief among these obstacles are the sensitivity and scalability of cell identification. The former is integral for enabling accurate single-cell analysis of more than the most ubiquitous cell types, while the latter becomes paramount when considering economics and the volume of samples necessary for validation of cellular biomarkers established via single-cell profiling. Standard methods in use today include variations and iterative improvements in traditional imaging and flow cytometry solutions^8,9^. However, the sensitivity and scalability of these methods are fundamentally limited by data noise and operational requirements^10,11^.

To address these issues, we have developed a biocytometry system that enables massively parallel multiparametric identification of targeted immunophenotypes with a single-cell sensitivity. The system employs *S. cerevisiae* S288C cells, engineered to specifically adhere to designated surface markers, facilitate local intercellular communication, and initiate reporter response upon detection of the targeted immunophenotype. By avoiding the limitations of traditional instruments and molecular probes, it adopts a streamlined workflow that can be integrated seamlessly with established analytical pipelines (Fig. 1a-c). The standard workflow implements basic mixing and incubation steps combined with high-throughput spectrophotometry to easily accommodate the simultaneous measurement of a wide range of immunophenotypes. The method has been validated on a diverse set of immortalized cell lines, HeLa, HaCaT, KG1a, and Jurkat, which exhibit a diverse set of surface markers, including EGFR, EpCAM, CD34, CD25, and phosphatidylserine. Targeted cell types were identified through a combination of surface markers that fulfilled logic statements, including AND, OR, and YES gates. The sensitivity of the standard workflow was validated in a controlled setting, and the limit of detection was 1 cell per million.

**Fig. 1.**
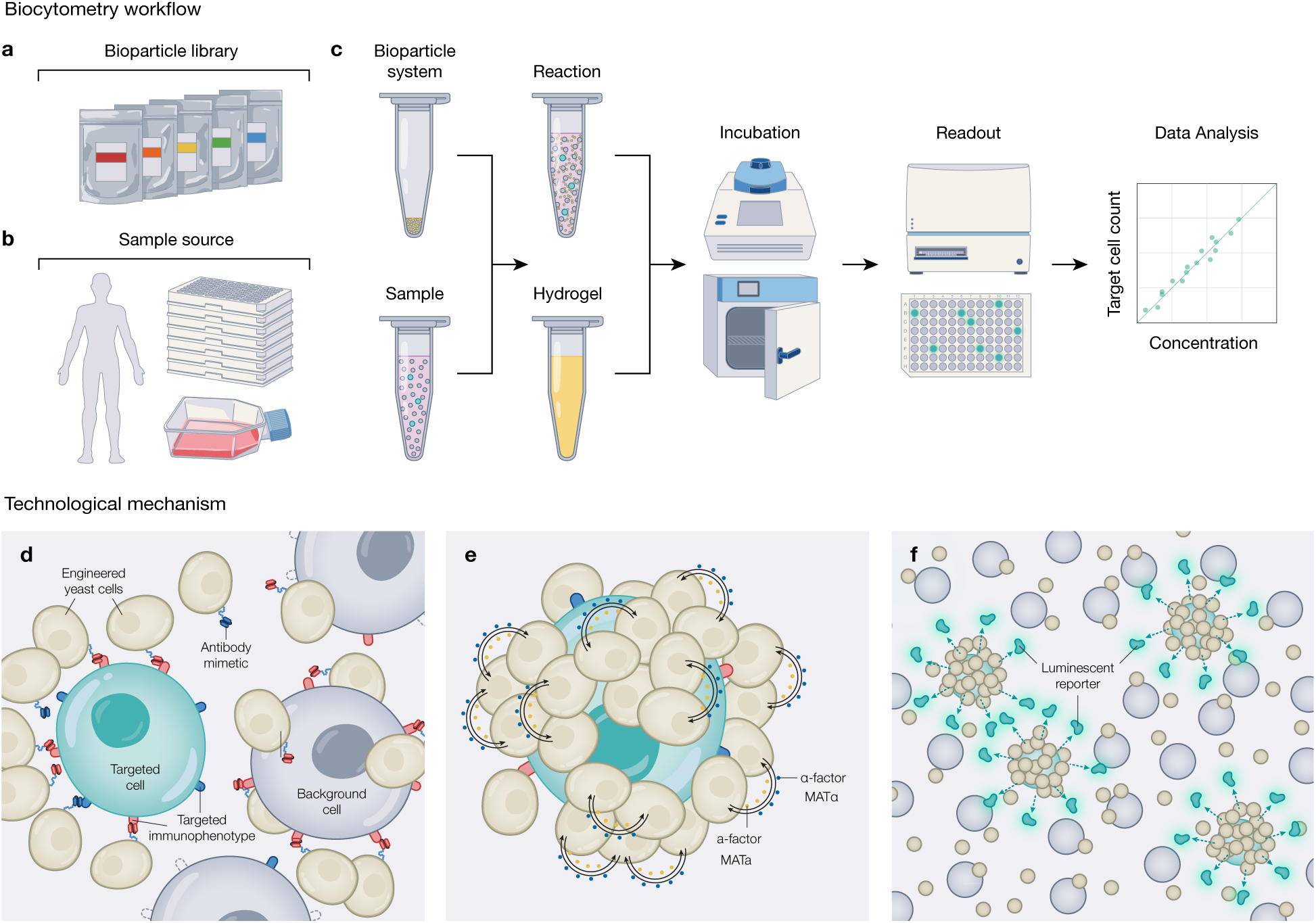
| Biocytometry general workflow and technological principle. (**a**) The platform workflow. A bioparticle system is designed to identify a given cell immunophenotype using a combination of bioparticle types, each defined by distinct binding and signaling properties. (**b**) The platform is robust to handling and to different sample types, including peripheral blood, cell lines, and cryopreserved or stabilized samples. (**c**) The process begins by combining the bioparticle system with a sample and hydrogel matrix. This is followed by an incubation period and finally a readout that determines the target cell count. (**d**) The technological mechanism. Bioparticles display antibody mimetics on their surface, defined by their binding type. These bioparticles effciently bind to cells in suspension, guided by each cell’s unique immunophenotype. (**e**) Within each bioparticle-cell complex, bioparticles secrete trace amounts of pheromone according to their signaling type. Target immunophenotypes yield positive feedback signaling patterns that reinforce signaling activity and ultimately switch bioparticles to an activated state. (**f**) Bioparticle activation is marked by the turning on of programmable reporter gene circuits, e.g., a luminescent reporter is secreted into the medium for the sensitive enumeration of target cells.

A comparative analysis of biocytometry and flow cytometry was performed on widely used clinical sample matrices, including peripheral blood, cryopreserved peripheral blood mononuclear cells (PBMCs), and primary cell cultures. Biocytometry was equivalent to flow cytometry in lower-sensitivity quantifications (>10^−4^ sensitivity) and outperformed flow cytometry in higher-sensitivity quantifications (<10^−4^ sensitivity), achieving the theoretical limit of detection. Moreover, the sensitivity and specificity remained undiminished when applied to challenging sample matrices (ROC, AUC = 0.996-1.000), notably apoptotic primary cell cultures and bone marrow. These results suggest that our technology may prove particularly effective in applications that demand regular high-sensitivity detection, such as the enumeration of hematopoietic stem cells^12^, CAR-T cells^13^, or distinct immune cell clonotypes^14^.

## Results

### Engineering of the biocytometry system

Biocytometry employs *S. cerevisiae* S288C cells (*MATa/MATα; ura3-52; trp1-289; leu2-3, 112; his3:1; MAL2-8C; SUC2*)^15^, engineered to initiate reporter response upon adherence and detection of targeted immunophenotype. The specificity toward targeted immunophenotypes is determined by the combination of bioparticle types, each of which is defined by distinct binding and signaling properties and is configurable for surface marker logical gating: AND, OR, YES. For instance, a bioparticle combination denoted by BIOS(EpCAM) operates under a YES gate to selectively identify EpCAM+ cells, while BIOS(EpCAM+EGFR) enables the identification of EpCAM+EGFR+ target cells. Up to 4 bioparticle types and a total of 10 or more bioparticles may contribute to a single identification event (Supplementary Fig. 1).

Specifically, the engineered bioparticles include (i) a novel anchor system that enables efficient formation of bioparticle-cell complexes (Fig. 1d); (ii) a hypersensitized signaling pathway to ensure robust inter-bioparticle communication and accurate interpretation of cellular immunophenotypes (Fig. 1e); and (iii) a reporter module that relays the output of logical processing to a luminescent reporter gene (Fig. 1f). Furthermore, (iv) a semipermeable hydrogel matrix isolates inter-bioparticle communication, allowing single-cell resolution without sample compartmentalization.

Antibody mimetics displayed on the surface of bioparticles enable the formation of immunospecific bioparticle-cell complexes. Anchor domains in existing yeast display systems, although extensively used for antibody engineering and protein-protein interaction studies^16–19^, exhibit limited efficacy in complex formation (Supplementary Fig. 2a). We engineered a novel cell wall anchor domain using a truncated version of yeast FLO11, spanning from the central region to the GPI anchor (residues 206-1367). Comparative assessments with HaCaT cells revealed a 652% increase in binding efficiency for bioparticles using our anchor system over using the conventional SAG1 anchor (Supplementary Fig. 2b). Notably, our system enables the seamless integration of established proteinaceous binding moieties, including but not limited to antibody conjugates, scFvs, DARPins, and Affibodies (Supplementary Fig. 2c).

A redirected mating pathway governs inter-bioparticle communication and facilitates the accurate immunophenotype interpretation of analyzed cells. Mating pathway activation in the bioparticle relies on the adjacency of the cognate signaling partners in the bioparticle-cell complex. For the scope of this article, we utilized yeast bioparticles of two mating types, MATa and MATα, that can produce mating factors (MFA1, MFX1) and detect them through GPCR receptors (STE2 and STE3)^20^. The strains were engineered for enhanced mating factor sensitivity and tunable production through the implementation of mating-type-specific alterations. MATα-origin bioparticles incorporate the SST2 protein under the regulation of a weaker *RAD27p* promoter (Supplementary Table 1). For bioparticles of MATa origin, increased sensitivity is realized solely through the removal of the BAR1 protease, circumventing any further modifications to SST2 expression. To make the mating factor production tunable, we deleted wild-type mating factors from the chromosome and integrated an octet of the corresponding mating factors under the control of the mating-factor-inducible *FIG1p* promoter (Supplementary Table 1). Our findings reveal a pronounced increase in mating pheromone sensitivity, with a 29.5-fold improvement on average, and in the ratio of active to basal pheromone production, with a 7.6-fold enhancement on average, both measured against their WT parent strains (Supplementary Fig. 3).

The reporter system provides rapid and sensitive identification of activated bioparticles. The detection of a single activated bioparticle-cell complex necessitates a system with exceptional induction fold change, high yield, and rapid expression kinetics. In the current work, we chose to implement NanoLuc luciferase under the control of an extended *FIG1(l)p* promoter (Supplementary Table 1). Incorporating an additional 70 bp upstream of *FIG1p* significantly enhanced promoter fold-induction properties by reducing basal expression 5-fold and increasing maximal expression levels 1.5-fold (Supplementary Fig. 4). Additionally, by altering the GPA1 subunit (Gα), we successfully dampened the basal activity of the mating pathway, leading to a 242% increase in reporter gene fold-induction (Supplementary Fig. 4). Collectively, these alterations resulted in at least a 10-fold enhancement in reporter gene fold-induction relative to the wild-type strain using a *FIG1p* regulated reporter (Supplementary Fig. 4). The system’s modular architecture also permits the integration of alternative reporters such as eGFP^21^ or β-lactamase^22^.

A functionalized semipermeable hydrogel matrix is crucial for confining communication to bioparticle-cell complexes and minimizing intercomplex cross-talk. Conventional single-cell analysis frequently necessitates sample compartmentalization, utilizing either drop-based or spatiotemporal isolation techniques^23,24^. To circumvent these limitations, biocytometry employs a hydrogel system formulated from a blend of biopolymers that enables free nutrient diffusion while accentuating local gradients of mating pheromones and actively suppressing their dispersion (Supplementary Fig. 5a). Additionally, bioparticles expressing BAR1 protease are incorporated into the hydrogel system, acting as concentration sinks to regulate the overall concentration of mating pheromones (Supplementary Fig. 5b). The synergistic interplay between the hydrogel matrix and the hypersensitized signaling pathway is pivotal for the fundamental functionality of the biocytometry system (Supplementary Fig. 5c).

### Technical validation with cell lines

Biocytometry’s capabilities in the accurate enumeration of targeted cell immunophenotypes were demonstrated with the HaCaT cell line, an immortalized keratinocyte model characterized by the expression of EpCAM and EGFR membrane proteins (Supplementary Fig. 6a). Comparable results were obtained for a broad collection of cell lines, such as HeLa, Jurkat, RPMI8226 and KG1a, characterized by unique sets of surface markers (Supplementary Fig. 6b).

Target cells were labeled with ATTO425-maleimide (Fig. 2a) and resuspended in 100 µl of PBS + 0.1% gelatin at quantities ranging from 0 to ∼1000 cells/sample. Analysis was performed using BIOS(EpCAM), in accordance with the biocytometry workflow. Luminescence readout was conducted using an optical bottom 96-well plate, followed by microscopic examination to facilitate the absolute quantification of target cells for subsequent data analysis (Fig. 2b).

**Fig. 2.**
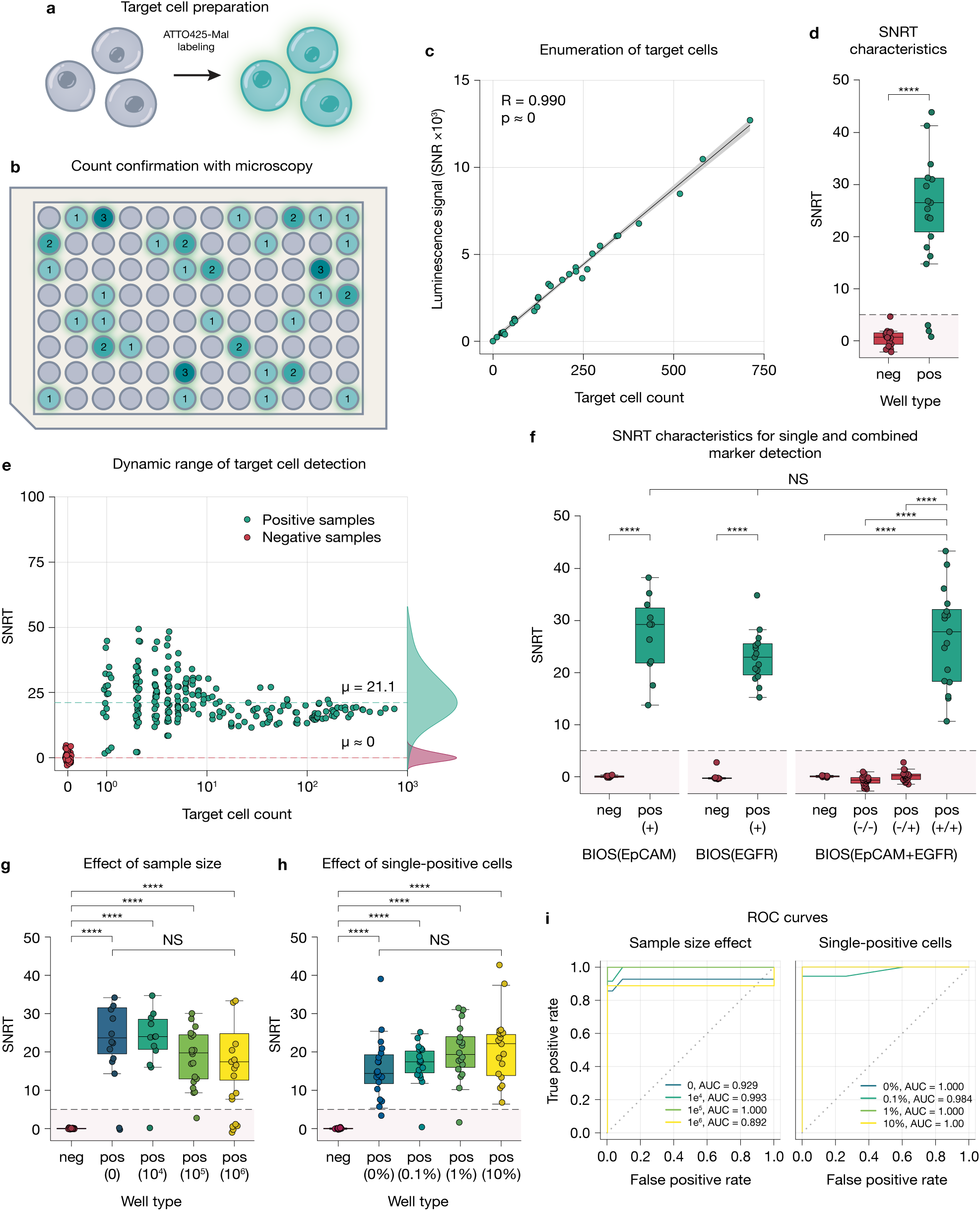
| Evaluation and benchmarking of sensitivity, dynamic range and multimarker detection. (**a**) Target cells labeled with ATTO425-maleimide enable post-analysis count confirmation via microscopy. (**b**) Combining well luminescence with microscopic counts yields the signal-to-noise ratio per target (SNRT), a measure of detection performance. (**c**) Target cell spiking experiment. Sample tubes were spiked with 0-1000 HaCaT (EpCAM+) target cells and processed according to the standard protocol with BIOS(EpCAM) bioparticles. The luminescence readout exhibited a strong correlation with the number of spiked targets (R = 0.990, p ≈ 0). The 95% confidence intervals are also shown (gray area). (**d**) Wells devoid of target cells consistently remained within 3 standard deviations of the blank wells, with the maximum signal-to-noise ratio (SNR) per well reaching 4.7, attesting to the inherent specificity of the method. A mean SNRT for wells containing a single target cell was established at 27.1. (**e**) The assay exhibits exceptional sensitivity, accurately identifying samples containing even a single target cell and maintaining consistent performance across a wide dynamic range (1 to 1,000 target cells/reaction). (**f**) The evaluation of the assay’s performance in detecting target cells using a dual-marker (EpCAM and EGFR) strategy is depicted. Sample tubes containing 0-30 target cells derived from HL-60 (EpCAM−EGFR−), HeLa (EpCAM−EGFR+), or HaCaT (EpCAM+EGFR+) cell lines were analyzed using either BIOS(EpCAM), BIOS(EGFR), or BIOS(EpCAM+EGFR) per the standard protocol. The mean SNRT values obtained were 27.0 for BIOS(EpCAM), 23.0 for BIOS(EGFR), and 26.7 for BIOS(EpCAM+EGFR), demonstrating comparable detection efficacy (ANOVA, p = 0.298). Signals from samples with zero-count target cells, HL-60 (double-negative), or HeLa (single-positive) target cells consistently fell beneath the threshold for positive classification. This underscores the platform’s discriminative ability in a multi-marker detection context, as confirmed by the results of Mann-Whitney U tests (zero-count: p = 2.11×10^−5^, HL-60: p = 1.26×10^−7^, HeLa: p = 1.71×10^−7^). (**g**) To assess the robustness of biocytometry across varying sample sizes, we integrated a fixed number (∼30 cells/sample) of EpCAM+ HaCaT target cells into HL-60 background cells (EpCAM−, 0 to 1×10⁶/sample). The BIOS(EpCAM) biocytometry system was utilized for analysis. Mean SNRT values remained stable across increasing background cell counts, ranging from 31.3 to 24.2, as confirmed by statistical analyses (ANOVA, p = 0.137; Mann-Whitney U, p < 10⁻⁶). HL-60 cells alone yielded no detectable signal. (**h**) To examine the influence of single-marker-positive cells on dual-marker detection, samples were prepared with a constant count of HaCaT target cells (EpCAM+EGFR+, ∼30 cells/sample) and varying numbers of HeLa cells (EpCAM−EGFR+, 0 to 1×10⁴/sample), all against a uniform HL-60 cell background (EpCAM−EGFR−, 1×10⁵ cells/sample). When BIOS(EpCAM+EGFR) was used according to the standard protocol, the assay maintained stable performance across differing HeLa cell counts. Mean SNRT values ranged from 16.7 to 20.6 (ANOVA, p = 0.280; Mann-Whitney U test, p < 10^−6^). Crucially, the assay retained specificity, as samples containing solely HeLa cells did not produce signals. (**i**) ROC curves were generated for each sample, both by background cell size—0, 1×10⁴, 1×10⁵, 1×10⁶ cells—and by the number of single-positive cells—0, 1×10², 1×10³, 1×10⁴. The corresponding AUC values were 0.939, 0.997, 1.000, and 0.917 for the background cell sizes and 0.939, 0.997, 1.000, and 0.917 for the single-positive cell counts. Error bars represent 1 s.d. ****, p < 0.0001; NS, nonsignificant. A line at SNR = 5 delineates the upper limit for negative samples.

The normalized luminescence values derived from the assay readout were plotted against the number of target cells within each tested sample (Fig. 2c). Biocytometry reproduced the microscopy results with high accuracy (R = 0.990, p ≈ 0). To further examine the acquired data, we conducted a well-level analysis with a single target cell (Fig. 2d). Wells devoid of target cells constantly remained within 3 standard deviations of the blank wells, with the maximum signal-to-noise ratio (SNR) per well reaching 4.7, attesting to the inherent specificity of the method. A mean SNR normalized by target cell count (SNRT) was established at 27.1.

To evaluate the dynamic range, we processed a series of samples containing varying numbers of HaCaT target cells (EpCAM+, ranging from 0 to 1×10^3^/sample) using the BIOS(EpCAM) and biocytometry workflow. Statistical analysis revealed consistent assay performance across the range of target cell counts (ANOVA, p = 0.930), with a mean SNRT value of 21.1. This value was subsequently utilized as the predictive parameter for target cell number estimations. The signal in wells devoid of target cells remained stable, exhibiting an average SNR ≈ 0 (Fig. 2e).

### Multiparametric target cell identification

While the identification of specific cellular types can be achieved using a single surface marker^25–27^, a more comprehensive identification typically requires multimarker analysis^28,29^. For multimarker analysis, EpCAM and EGFR were used as the target surface markers. Prior to the combined utilization of EpCAM and EGFR, we performed an experiment to verify the absence of marker bias within our platform (Fig. 2f, left). Following the established protocol, target cell quantification was performed with fluorescently labeled HaCaT cells (EpCAM+EGFR+, ∼30 cells/sample), utilizing either BIOS(EpCAM) or BIOS(EGFR). The data demonstrated no statistically significant difference in the performance of the two marker-specific bioparticles (two-sided t test, p = 0.114). Both BIOS(EpCAM) and BIOS(EGFR) supported robust detection of the target cells (Mann-Whitney U test, p < 10^−4^), with mean SNRT values of 27.0 and 23.0, respectively.

Having established equivalent performance for BIOS(EpCAM) and BIOS(EGFR) bioparticle systems, we applied them simultaneously in dual-marker detection assays (Fig. 2f, right). For this purpose, we assessed target cells (∼30 cells/sample) from various cell lines: HaCaT (EpCAM+EGFR+), HeLa (EpCAM−EGFR+), and HL-60 (EpCAM−EGFR−), employing BIOS(EpCAM+EGFR) in the biocytometry workflow. Utilizing the two markers in combination yielded a detection performance on par (SNRT = 26.7; Mann-Whitney U test, p < 10^−4^) with the use of either marker individually (ANOVA, p = 0.298). Critically, cells devoid of expression for both markers remained under the threshold for positive classification.

### Limits of target cell identification in mixed samples

In the previous experiment, single-target sensitivity was established. To evaluate the upper limit of sample size amenable to single-cell resolution, we conducted an experiment using the HL-60 cell line^30^, a suitable proxy for the leukocyte fraction commonly examined in clinical settings. We introduced a constant number of HaCaT target cells (EpCAM+, ∼30 cells/sample) while varying the quantity of HL-60 background cells (EpCAM−, 0 to 1×10^6^/sample) and processed these samples using BIOS(EpCAM) following the biocytometry workflow. The data suggest that assay performance remains stable across varying background cell concentrations (ANOVA, p = 0.137; Mann-Whitney U test, p < 10^−6^), with mean SNRT values of 31.3, 28.3, 22.0, and 24.2 for samples with 0, 1×10^4^, 1×10^5^, and 1×10^6^ background cells, respectively (Fig. 2g). Notably, HL-60 cells on their own did not produce any discernible signal (max SNR = 0.3). The assay exhibited high sensitivity, with AUC values of 0.929, 0.993, 1.000, and 0.892 recorded for samples containing 0, 1×10^4^, 1×10^5^, and 1×10^6^ background cells, respectively (Fig. 2i, left). With a sensitivity allowing the identification of one target cell among 1×10^6^ total cells, biocytometry could substantially enhance the routine detection of rare cellular subpopulations, such as hematopoietic stem cells^12^, CAR-T cells^13^, or distinct CD4+ or CD8+ T-cell clonotypes^14^, beyond the conventional limit of 1×10^−4^ set by flow cytometry^31^.

To evaluate the influence of single-marker positive subpopulations on the resolution of dual-marker positive target cells, HeLa cells were introduced into the HL-60 background. Samples were prepared with a fixed count of HaCaT target cells (EpCAM+EGFR+, ∼30 cells/sample) while varying the number of HeLa cells (EpCAM−EGFR+, 0 to 1×10^4^/sample), all against a constant background of HL-60 cells (EpCAM−EGFR−, 1×10^5^ cells). The samples were then processed as per our standard protocol using BIOS(EpCAM+EGFR). No change in assay performance was observed (ANOVA, p = 0.280; Mann-Whitney U test, p < 10^−7^) despite the increased count of single-positive cells (Fig. 2h). Mean SNRT values of 16.7, 17.4, 20.0, and 20.6 were obtained for samples containing 0, 1×10^2^, 1×10^3^, and 1×10^4^ single-positive cells, respectively. Crucially, the presence of HeLa cells did not yield nonspecific signals, resulting in a maximum SNR of 0.4, comparable to the HL-60 background sample maximum of 0.3. ROC analysis of the assay yielded AUC values of 1.000, 0.984, 1.000, and 1.000 across samples containing 0, 1×10^2^, 1×10^3^, and 1×10^4^ single-positive cells, respectively (Fig. 2i, right). Our results confirm the effective use of two distinct bioparticle types for primary and secondary markers, with assay performance remaining consistent even when partial target cell population constitutes up to 10% of the total sample, a fraction within observed ranges for immune cell subsets^32^.

### Comparative assessment across complex sample matrices

Biocytometry and flow cytometry were further compared in a rigorous study on a large set of clinically relevant sample matrices. A set of samples was prepared by introducing varying quantities of unlabeled HaCaT cells (EpCAM+, 0, 4, 16, 64, 256, 1024/sample) into a variety of sample matrices, which included peripheral blood, cryopreserved PBMCs, and primary cell cultures (Fig. 3a). Each sample contained the equivalent of 1×10^6^ leukocytes. The resulting samples were subsequently split and processed separately. Biocytometric analysis utilized BIOS(EpCAM), adhering to the biocytometry workflow. Concurrently, flow cytometry samples were stained with AF647-conjugated anti-EpCAM antibody, following the flow cytometry workflow. Upon analysis, a strong correlation was observed between the two methods in peripheral blood (R = 1.000, p ≈ 0 for biocytometry; 0.943, p = 0.0032 for flow cytometry), cryopreserved PBMCs (R = 1.000, p ≈ 0 for biocytometry; R = 1.000, p ≈ 0 for flow cytometry) and primary cell cultures (R = 1.000, p ≈ 0 for biocytometry; R = 1.000, p ≈ 0 for flow cytometry). For samples with lower target cell counts (0-64 spiked target cells, < 10^−4^ sensitivity), flow cytometry analysis was impeded by its limit of blank (LOB), failing to quantify below this level. Significant variability in flow cytometry LOB was observed across examined sample matrices, with cryopreserved PBMCs yielding the highest false-positive rate at 121 per million cells analyzed, followed by lysed blood at 55 per million and primary cell cultures at 31 per million. Biocytometry, on the other hand, maintained a null LOB across all tested sample matrices (Fig. 3b).

**Fig. 3.**
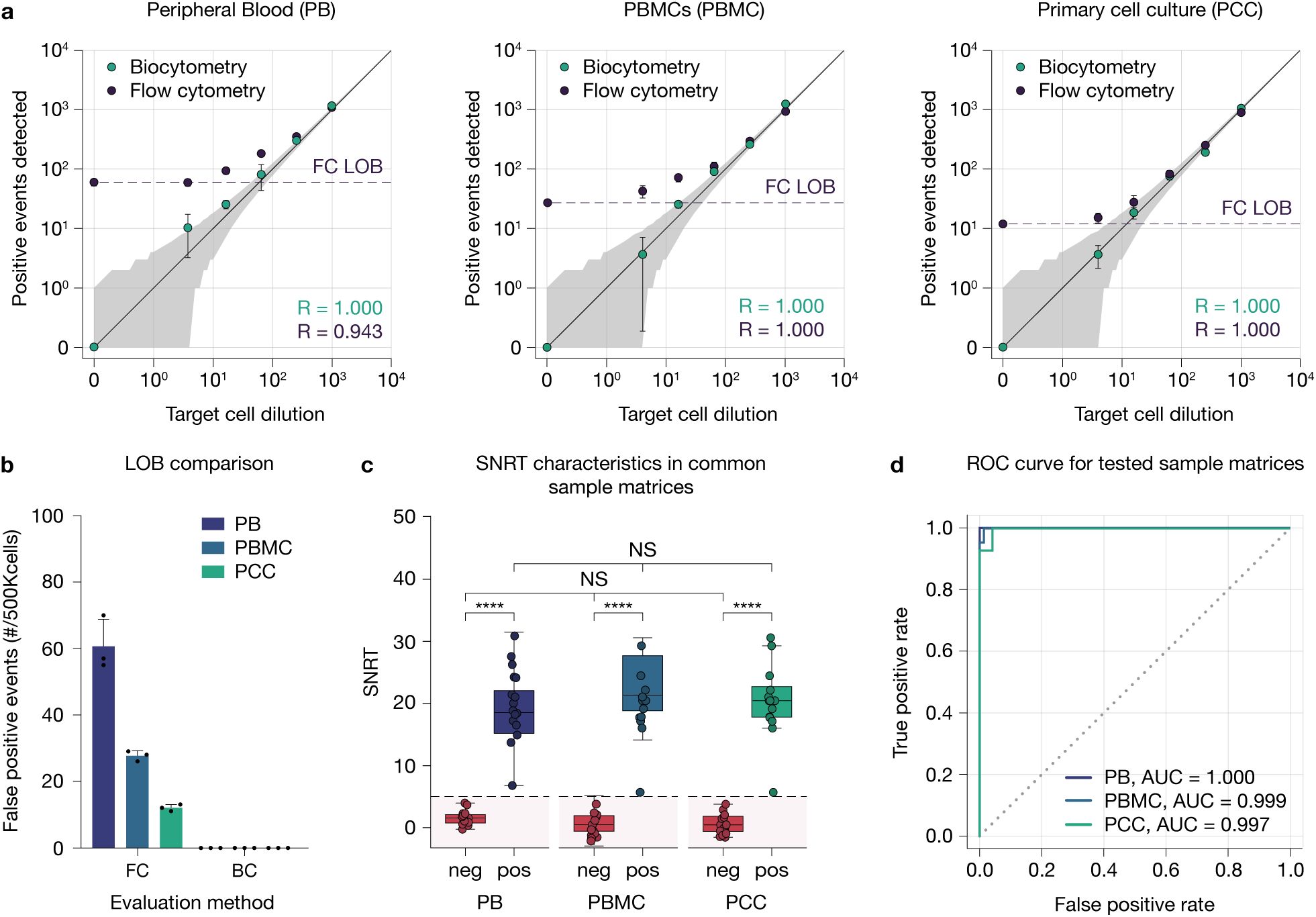
| Comparison of detection limits for the platform and flow cytometry in common sample matrices. (**a**) For a comparative analysis of the platform and flow cytometry, samples with different HaCaT (EpCAM+) target cell quantities (0, 4, 16, 64, 256, 1024) were prepared in a range of sample matrices, including peripheral blood (PB), PBMCs (PBMC), or primary cell culture (PCC), each containing approximately 5×10^5^ leukocytes. Samples (n = 3 independent replicates) in a target cell dilution series were analyzed in parallel. Our platform utilized BIOS(EpCAM), adhering to the biocytometry workflow. For flow cytometry, samples were stained with AF647-conjugated anti-EpCAM antibody, following the flow cytometry workflow. We observed a strong correlation between the two methods and the target cell dilution series for samples containing high target cell counts (64-1024 spiked target cells, >10^−4^ sensitivity). For samples with lower target cell counts (0-64 spiked target cells, <10^−4^ sensitivity), flow cytometry analysis was impeded by its limit of blank (FC LOB), failing to quantify below this level. Spearman’s (R) is shown. The uncertainty in target cell spiking counts is highlighted (gray area). (**b**) In contrast to flow cytometry (FC), which showed varying false-positive rates among tested sample matrices—highest in peripheral blood (121 per million cells analyzed), followed by PBMCs (55 per million), and primary cell cultures (31 per million) —biocytometry (BC) consistently exhibited a null limit of blank (LOB) across all sample types. (**c**) The assay’s performance in absolute quantification within complex sample matrices was also evaluated. Microscopic examination was employed to validate the exact count of the target cells. The mean SNRT values for target cell detection—17.9 for peripheral blood, 21.6 for PBMCs, and 19.3 for primary cell cultures—indicated consistent performance across all tested matrices (ANOVA, p = 0.991). Moreover, the assay exhibited robust detection characteristics, as demonstrated by the Mann-Whitney U test results (PB: p = 3.72×10^−6^, PBMC: p = 2.57×10^−5^, PCC: p = 1.76×10^−6^). (**d**) ROC curves were generated for each sample matrix, revealing AUC values of 1.000, 0.999, and 0.997 for peripheral blood, PBMCs, and primary cell cultures, respectively. Error bars represent 1 s.d. ****, p < 0.0001; NS, nonsignificant. A line at SNR = 5 delineates the upper limit for negative samples.

To validate the platform’s capacity for absolute quantification in complex sample matrices, we introduced a constant number of fluorescently labeled HaCaT target cells (EpCAM+, ∼30 cells/sample) into a variety of sample matrices, including peripheral blood, cryopreserved PBMCs, and primary cell cultures. Each sample contained the equivalent of 5×10^5^ leukocytes. The analysis was performed with BIOS(EpCAM), adhering to the biocytometry workflow. Microscopic examination was performed to confirm the exact count of HaCaT cells in a sample. The assay’s performance was uniform (ANOVA, p = 0.991; Mann-Whitney U test, p < 10^−4^) in all evaluated matrices, as signified by mean SNRT values of 17.9, 21.6, and 19.3 for peripheral blood, cryopreserved PBMCs, and primary cell cultures, respectively (Fig. 3c). Consistency across different matrices was further validated by the ROC curves, which displayed AUC values of 1.000 for peripheral blood, 0.999 for cryopreserved PBMCs, and 0.997 for primary cell cultures (Fig. 3d).

### Platform compatibility with challenging sample matrices

Biocytometry was evaluated in the analysis of primary cell cultures exhibiting a high fraction of apoptotic cells. Apoptotic cells, arising from various factors such as sample age^33^, treatment^34^, or manipulation^35^, present analytical challenges in the analysis. Apoptosis in primary cell cultures was induced via brief heat shock^36^. The apoptotic rate of the cell cultures rose significantly, with 11.3% of cells identified as apoptotic (Fig. 4a), as confirmed by fluorescence staining using Annexin V FITC conjugate (Fig. 4b). A set of samples was prepared by introducing varying quantities of unlabeled HaCaT cells (EpCAM+, 0, 4, 16, 64, 256, 1024/sample) into a suspension of apoptotic primary cells. Each sample contained the equivalent of 1×10^6^ leukocytes. The resulting samples were subsequently split and processed separately. Biocytometric analysis was performed with BIOS(EpCAM), adhering to the biocytometry workflow. Concurrently, flow cytometry samples were stained with an AF647-conjugated anti-EpCAM antibody, following the flow cytometry workflow. Flow cytometry analysis revealed a marked 358% escalation in false-positive events, with a corresponding LOB increase to 111 false-positive events per million cells analyzed (Fig. 4c). In stark contrast, biocytometry maintained a null LOB, consistent with prior findings. Linear regression demonstrated a strong correlation for biocytometry (R = 1.000, p ≈ 0), in contrast to the suboptimal performance of flow cytometry (R = 0.867, p = 0.038) (Fig. 4d). During the identification of fluorescently labeled HaCaT cells, biocytometry exhibited nominal performance (Mann-Whitney U test, p < 10^−4^), registering a mean SNRT of 25.6 and an AUC of 1.000 in ROC analysis (Fig. 4e and h).

**Fig. 4.**
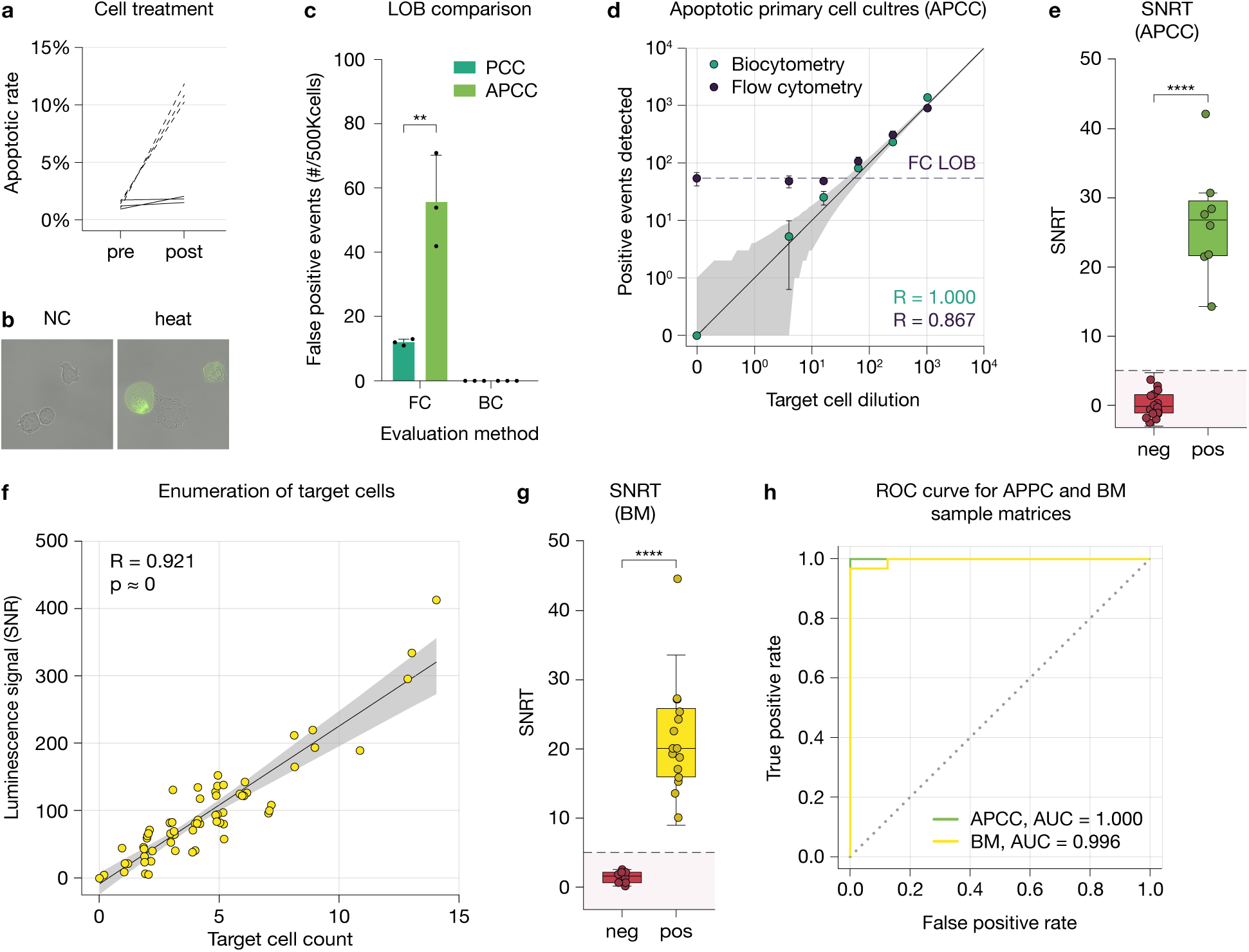
| Performance of biocytometry is not affected by challenging sample matrices. (**a**) The apoptotic rate in primary cell cultures increased after heat shock treatment, with 11.3% of cells confirmed as apoptotic. (**b**) Annexin V-FITC staining highlights the morphological features of apoptotic cells in microscopy images. (**c**) Biocytometry (BC) maintained a null LOB, in contrast to the escalated LOB observed with flow cytometry (FC). (**d**) A scatter plot analysis reveals a strong correlation for biocytometry (R = 1.000, p ≈ 0) and suboptimal performance for flow cytometry (R = 0.867, p = 0.038) in apoptotic primary cell cultures. Spearman’s (R) is shown. Uncertainty in target cell spiking counts is highlighted (gray area). (**e**) With fluorescently labeled HaCaT cells used in apoptotic primary cell cultures (APCCs), biocytometry registers a mean SNRT of 25.6. (**f**) A strong correlation (R = 0.921, p ≈ 0) was observed between the estimated target cell counts in bone marrow and biocytometry readouts. The 95% confidence intervals are also shown (gray area). (**g**) The mean SNRT in bone marrow was 20.0, demonstrating comparable performance to other matrices. (**h**) ROC curve analysis yields an AUC value of 1.000 in APCC and 0.996 in bone marrow. Error bars represent 1 s.d. ****, p < 0.0001; NS, nonsignificant. A line at SNR = 5 delineates the upper limit for negative samples.

Biocytometry was evaluated for the analysis of bone marrow aspirates. Bone marrow is especially difficult to analyze due to the complex microenvironment^37^, debris^38,39^, and small sample size^40^. To explore whether biocytometry could mitigate these challenges, we spiked bone marrow samples from a patient undergoing hematopoietic stem cell transplantation (HSCT) for acute myeloid leukemia (AML) with fluorescently labeled HaCaT target cells (0-30 cells/sample) and analyzed them using BIOS(EpCAM). Given the limited sample, flow cytometry analysis was not conducted. The experimental data showed an unaltered baseline LOB, along with a significant correlation (R = 0.921, p ≈ 0) between target cell counts estimated by our platform and those confirmed through fluorescence microscopy (Fig. 4f). Furthermore, the target cell detection performance proved comparable to that in other sample matrices (Mann-Whitney U test, p < 10^−4^), as indicated by a mean SNRT of 20.0 (Fig. 4g). The ROC curve for the assay yielded an AUC value of 0.996 (Fig. 4h).

### Quantification of leukocyte activation in PBMCs

Biocytometry was evaluated for the quantification of leukocyte activation status. CD25 is a putative marker for long-term leukocyte activation. The extent of leukocyte activation in assays can range from low levels when evaluating particular T-cell receptor (TCR) subsets^41^ to nearly universal when assessing systemic immune function^42^. To obtain a high activation rate, we treated a PBMC sample from a healthy donor with the nonspecific mitogen PHA (2.5 µg/ml) alongside an uninduced control sample. To capture varying activation rates, we combined the induced and uninduced control samples at different ratios (0%, 11%, 33% and 100% induced). Each sample contained the equivalent of 1×10^6^ leukocytes. The resulting dilution series was evaluated in parallel by biocytometry and flow cytometry. In compliance with the biocytometry workflow, the biocytometry assay utilized BIOS(CD25), while flow cytometry was performed according to the flow cytometry protocol using FITC-conjugated anti-CD25 antibodies. A robust correlation was observed between biocytometry and flow cytometry for estimating CD25+ events, as indicated by a Spearman correlation coefficient of R = 0.975 (p < 10^−9^) (Fig. 5a). To assess the degree of agreement between the two methodologies, Bland-Altman analysis was performed. The resultant plot disclosed a mean bias of 0.25%, with limits of agreement spanning from −1.83% to 2.33% (Fig. 5b).

**Fig. 5.**
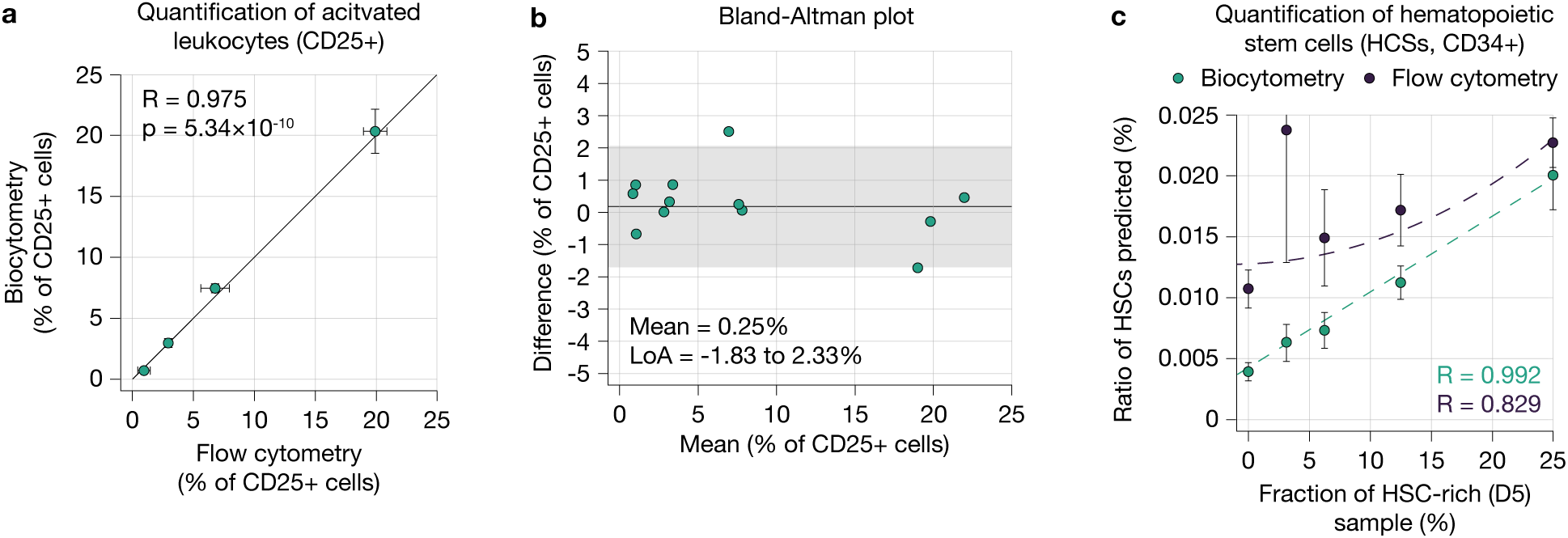
| Comparative analysis of biocytometry and flow cytometry in low-and high-sensitivity applications. (**a**) Correlation plot showing a strong relationship (R = 0.975, p < 10^−9^) between CD25+ events estimated by biocytometry using the BIOS(CD25) bioparticle system and flow cytometry with FITC-conjugated anti-CD25 antibodies. (**b**) Bland-Altman plot indicating a mean bias of 0.25% and limits of agreement from −1.83% to 2.33% between the two methods for leukocyte activation assessment. (**c**) Scatter plot illustrating the enumeration of CD34+ hematopoietic stem cells (HSCs) in a dilution series derived from D0 and D5 samples during G-CSF-induced mobilization. Biocytometry achieved a near perfect linear fit (R = 0.992, p ≈ 0), in contrast to the suboptimal performance of flow cytometry at low HSC concentrations. The CVs were 18.4% for biocytometry and 24.9% for flow cytometry, with flow cytometry displaying a higher maximum CV (45.2%) near the LOD.

### HSC enumeration in mobilized patients

Biocytometry was evaluated for the enumeration of CD34+ hematopoietic stem cells (HSCs), a critical indicator in stem cell mobilization procedures^43^. Patients receiving granulocyte colony-stimulating factor (G-CSF) for stem cell mobilization provided peripheral blood samples both at the onset of mobilization (D0) and at the time of leukapheresis (D5). PBMCs were isolated via conventional Ficoll density gradient separation. A dilution series was established using D0 and D5 samples, generating a range of samples with increasing HSC content. Each sample contained the equivalent of 1×10^6^ leukocytes. The samples were subjected to parallel analysis using a biocytometry workflow with BIOS(CD34) and flow cytometry following a quantification protocol using PE-conjugated anti-CD34 antibodies. The biocytometry results exhibited a strong linear relationship with the HSC dilution series (R = 0.992, p ≈ 0). Conversely, flow cytometry yielded poor correlation (R = 0.829, p ≈ 0), a limitation attributable to HSC concentrations nearing the detection limit (Fig. 5c). The two methods exhibited comparable coefficients of variation (CVs): 18.4% for biocytometry and 24.9% for flow cytometry. Flow cytometry showed a notably higher maximum CV (45.2%) than biocytometry (25.0%) near the limit of detection (LOD). The observed LOD aligns with the baseline HSC levels in healthy populations and is consistent with values reported in ISHAGE interlaboratory harmonization studies (Supplementary Fig. 7).

### Measurement of the cell-specific apoptosis rate

Biocytometry was evaluated for measuring cell-specific apoptosis in mixed cell cultures. Despite the growing trend toward studying apoptosis in complex biological systems such as organoids and mixed cell cultures^44,45^, conventional methods such as Caspase-based assays and MTT tests still yield limited sensitivity and quantify only aggregate apoptosis levels^46^. To assess cell-specific apoptosis, we utilized mixed cultures of HaCaT (HCT, EpCAM+) and Jurkat (JUR, EpCAM−) cells at varied ratios (1:0 to 1:8). Cells were cultured at a density of 40,000 cells/well in RPMI medium supplemented with 10% FBS and either subjected to 1 mM DTT treatment or left untreated. Following a 2.5-hour incubation at 37°C in a 5% CO_2_ atmosphere, both total and apoptotic HaCaT cell counts were established using BIOS(EpCAM) and BIOS(EpCAM+PS), adhering to the biocytometry protocol. A noticeable reduction in total EpCAM+cell count was evident in induced cell cultures, alongside an increase in the number of apoptotic EpCAM+ cells, aligning with the anticipated apoptotic changes (Fig. 6a). Apoptotic rates were calculated by normalizing BIOS(EpCAM+PS) counts to total cell counts from BIOS(EpCAM). We observed a 7.2-fold increase in apoptotic rates among EpCAM+ cells exposed to 1 mM DTT, reaching 12.2% compared to a 1.7% baseline in untreated controls (Fig. 6b). Importantly, the apoptotic rate remained consistent across varying Jurkat cell ratios, underscoring the assay’s specificity for HaCaT cells, which was further corroborated by the absence of measurable apoptotic events in HaCaT-devoid samples (Fig. 6c).

**Fig. 6.**
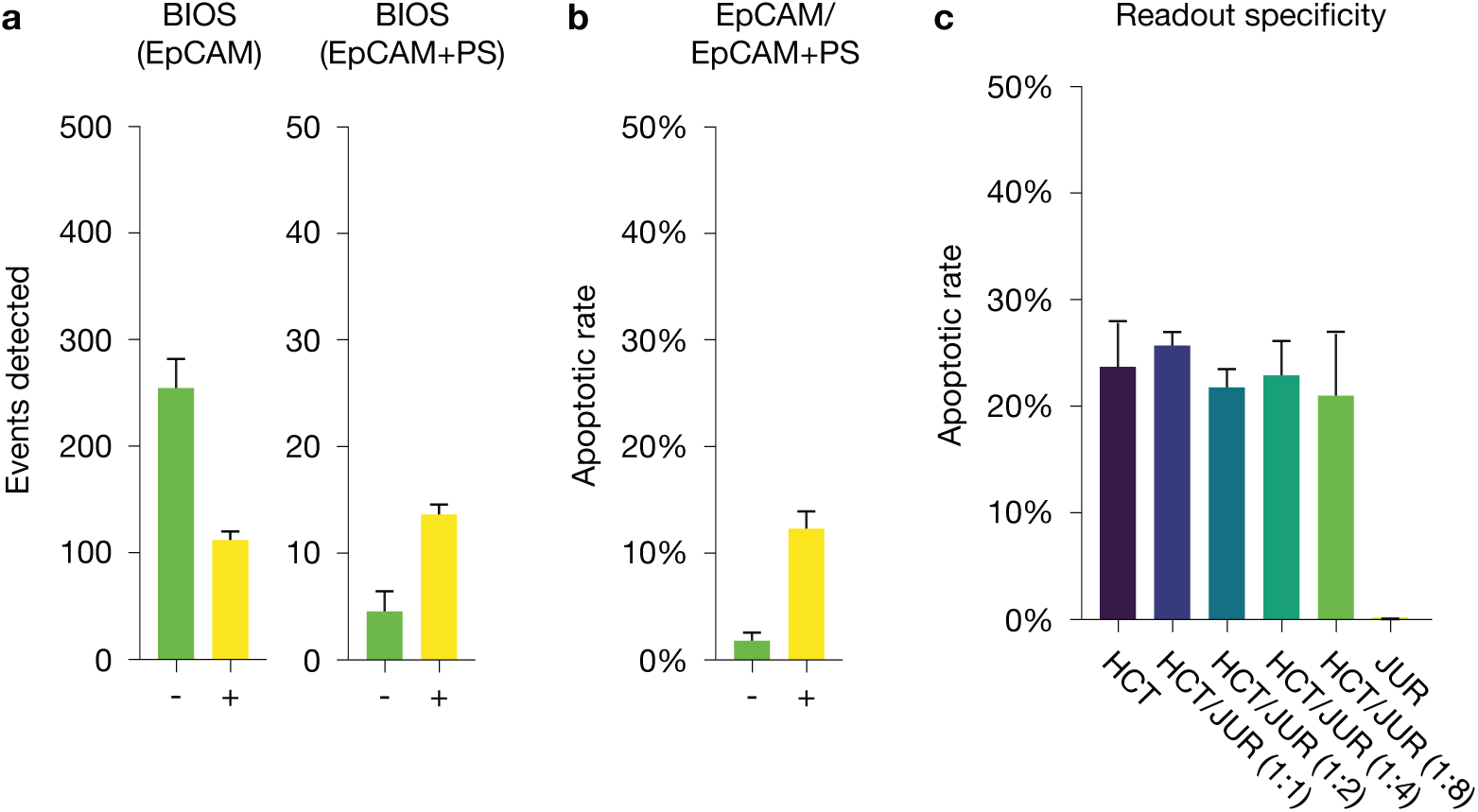
| Biocytometric quantification of apoptosis in mixed cell cultures. To assess the specificity of the apoptosis readout obtained via biocytometry, mixed cell cultures comprising HaCaT (HCT, EpCAM+) and Jurkat (JUR, EpCAM−) cells were employed at varying ratios (1:0, 1:1, 1:2, 1:4, 1:8, 0:1). Cells were cultured at a density of 40,000 cells/well in RPMI medium enriched with 10% FBS and either treated with 1 mM DTT or left untreated. Following a 2.5-hour incubation at 37°C in a 5% CO_2_ atmosphere, BIOS(EpCAM) was used to quantify the total number of HCT cells, whereas BIOS(EpCAM+PS) was used to assess the apoptotic status of HCT cells. (**a**) A noticeable decrease in the total count of EpCAM+ cells was observed, aligning with the anticipated apoptotic and subsequent necrotic changes in the cell culture, as corroborated by the BIOS(EpCAM+PS) readout. (**b**) Upon normalization of the signals from BIOS(EpCAM+PS) to BIOS(EpCAM), a robust response of HCT cells to DTT induction was evident. (**c**) The apoptotic rate remained consistent irrespective of the JUR (EpCAM−) cell count, confirming the specificity for detecting apoptosis in HCT (EpCAM+) cells, as evidenced by the absence of measurable events in HCT-devoid samples.

## Discussion

We report the development of biocytometry, a novel method for multiparametric immunophenotyping in suspension. The superior sensitivity and scalability of this method were demonstrated across a diverse set of sample types, ranging from 2D cell cultures to bone marrow aspirates, through the successful identification of 12 distinct cell types.

Biocytometry introduces a divergent analytical framework, a potential milestone, that enhances high-sensitivity cell analysis amidst ongoing reproducibility issues^47^. The marked improvements over traditional cell analysis methods, such as flow cytometry, are as follows: (i) biocytometry performs massively parallel integration of cell-surface marker profiles in vitro, allowing the simultaneous measurement of thousands of assays; (ii) biocytometry leverages cell-cell interactions over molecule-cell interactions, mitigating nonspecific binding and approaching the theoretical sensitivity limits^48,49^; and (iii) biocytometry utilizes a streamlined homogeneous assay workflow, ensuring highly sensitive and reproducible sample evaluation, which eliminates the necessity for specialized training and broadens the accessibility of cell analysis^50^.

The technical validation of biocytometry revealed strong alignment with established technologies over a wide dynamic range while supporting multiparametric identification down to a single cell within large sample sizes. The validation experiments utilized HaCaT cells, an immortalized keratinocyte model characterized by the expression of EpCAM and EGFR surface markers that is critical to both diagnostic and prognostic applications^51–53^. Cell enumeration assays yielded cell counts in concordance with microscopy observations (R = 0.990). Biocytometry accurately identified samples containing as little as a single target cell and maintained consistent performance across a wide dynamic range (1 to 1,000 target cells/sample). Multiparametric identification via an AND gate for EpCAM and EGFR surface markers successfully detected HaCaT cells (EpCAM+EGFR+), while control lines HeLa (EpCAM−EGFR+) and HL-60 (EpCAM−EGFR−) generated signals indistinguishable from those of negative samples. The sensitivity threshold for routine assessment was established at 1×10^−6^, well beyond the conventional limit of 1×10^−4^ set by flow cytometry^31^. Notably, the efficacy of multiparametric identification was uncompromised by the presence of cells with partial target immunophenotypes, even when they constituted up to 10% of the sample.

In a direct comparison to conventional flow cytometry, biocytometry demonstrated compatibility with clinically relevant sample types, consistently achieving superior sensitivity and specificity. The comparative study utilized HaCaT cells spiked into various sample matrices: peripheral blood, cryopreserved PBMCs, and primary cell cultures. For samples containing high numbers of target cells (128-1024 cells/million total cells; >10^−4^ sensitivity), both methods showed strong correlation across all matrices (R = 0.943-1.000). However, at lower target cell concentrations (0-64 cells/million total cells; <10^−4^ sensitivity), flow cytometry analysis was constrained by its LOB. Furthermore, these limitations varied among different sample matrices, which impedes the scalability of flow cytometry, as corroborated by previous studies^39^. In contrast, biocytometry reliably maintained a null LOB in all sample matrices, providing a robust framework for monitoring of cellular biomarkers, which is crucial for understanding disease dynamics and tailoring more precise treatments^54,55^.

The versatility of biocytometry was further highlighted when applied to analysis in notoriously challenging sample matrices such as apoptotic primary cell cultures and bone marrow aspirates, complicated by high levels of stromal components and debris and by limited sample availability^56,57^. In the context of apoptotic samples, flow cytometry showed a significant LOB increase of 358%, whereas biocytometry exhibited no such vulnerability, maintaining nominal target cell quantification metrics (R = 1.000). Likewise, for bone marrow samples, biocytometry achieved a near-perfect AUC of 0.996 in ROC analysis. Reliable performance in bone marrow samples is important for the applicability in MRD assessment, especially in the context of limited sample availability due to frequent dry tap occurrences in patients receiving multiple ablation therapies^58^.

Biocytometry was evaluated in three exemplary application domains: the quantification of leukocyte activation, enumeration of HSC populations, and measurement of cell-specific apoptosis. In quantifying activated leukocytes, biocytometry aligned closely with traditional flow cytometry, as evidenced by a Bland-Altman plot with a mean bias of 0.25% and limits of agreement between −1.83% and 2.33%. In HSC enumeration, biocytometry exhibited a strong linear relationship with the dilution series (R = 0.992), outperforming flow cytometry, which demonstrated a weaker correlation (R = 0.829), consistent with ISHAGE interlaboratory harmonization studies (Supplementary Fig. 7). Through multiparametric identification, we were able to discern the apoptotic rate of HaCaT cells in mixed cultures with Jurkat cells, marking the first reported use of a homogeneous assay workflow for targeted apoptosis quantification.

Through significant improvements across multiple performance metrics, biocytometry is in a unique position to reshape key sectors in diagnostics and therapy. Specifically, biocytometry, along with its imminent extensions, is well suited for facilitating the monitoring of rare premalignant subpopulations in conditions such as SMM^59^, enhancing the detection of circulating tumor cells (CTCs)^60^, advancing pharmacokinetic assessments in CAR-T therapeutic regimens (post-infusion/contraction and cellular subsets)^13^, and improving the accuracy of MRD evaluations^61^. Furthermore, biocytometry can facilitate the detection of rare clonotypes expressing unique TCR and BCR repertoires, making vaccine efficacy studies more reliable^15^. Biocytometry can also significantly increase the throughput of cell-specific readouts for in vitro drug discovery using more human-like samples, a high priority area enabling a shift from animal models^62–64^.

This study primarily contrasts biocytometry with flow cytometry, the established benchmark in rare cell analysis. However, alternative methods exist. Methods leveraging physical attributes such as cell size^65^, density^66^, or deformability^67^ offer the advantage of marker-independent detection in specific contexts and can achieve a substantial target cell recovery (approximately 75% capture rate for cells larger than 15 µm using a microsieve)^68^. Nevertheless, they exhibit reduced specificity compared to surface-marker-based approaches, often mandating subsequent analysis through low-throughput imaging techniques. Conversely, immunocapture-based approaches, particularly microfluidic technologies that combine physical separation with antibody recognition (e.g., nanotube CTC-Chip), frequently achieve exceptionally high purities in targeted cell populations and can facilitate downstream processing^69^. However, their scalability is limited by issues such as clogging, the need for bulky instrumentation, and significant operational costs^70^.

While this proof-of-concept study addresses what we believe to be the most critical limitations of cell analysis today, i.e., sensitivity and scalability, other aspects of the technology may prove to be equally valuable. For instance, biocytometry is not limited in target size and extends naturally to the quantification of cell-cell interactions, spheroids, and other irregular objects. Biocytometry may also enable the measurement of other data types that are inaccessible by traditional methods. Future enhancements to biocytometry are planned, including the implementation of logic gates for three or more markers, accommodation of larger sample sizes (up to 10^8^), and support for multiparametric data acquisition. It is our belief that biocytometry, coupled with new single-cell datasets, will soon lead to translational products in diagnostics, manufacturing, and drug discovery and will ultimately enable precision medicine at scale.

## Online methods

### Assessment of binding efficiency

Target cells were resuspended at a concentration of 300 cells per reaction in 100 µl of PBS containing 0.1% gelatin (Sigma-Aldrich, G9391). These samples were then subjected to the outlined biocytometry workflow (elaborated in subsequent sections) until diverging at the resuspension stage, where instead of a hydrogel matrix, the reaction mixture was distributed into a 96-well plate to facilitate microscopic analysis of cell-bioparticle complexes. A representative subset of cell-bioparticle complexes (n = 20) was randomly selected for each anchor system and evaluated. The samples were examined using microscopy (Olympus iX73 microscope, ×10 objective, BF channel). The captured images were analyzed via ImageJ software, utilizing a defined thresholding and particle counting protocol to quantify the number of bioparticles per complex. Statistical analysis was performed to ascertain the binding efficiency, employing t tests, to compare across different anchor systems and ascertain the significance of observed differences.

### Mating pheromone sensitivity quantification

Bioparticles were suspended in yeast extract-peptide-dextrose (YPD) medium to achieve an optical density (OD) of 0.1, with 100 µl aliquots placed in a 96-well plate. An initial dose of mating factor (IDT, custom peptide manufacturing) was added to the first well to reach a final concentration of 10 µM. A serial dilution was conducted across the subsequent wells by transferring 20 µl from one well to the next. The plate was incubated for 1 hour at 30°C in a shaking incubator (BioTek Synergy H1, 300 RPM). For readout, 11.4 ml of 0.1 M Tris (pH 8.6) was mixed with 300 µl of 1 µM furimazine (Promega, N1110) stock and 300 µl of 0.1% cycloheximide stock (Sigma-Aldrich, 01810). After incubation, the readout buffer was introduced to each well to facilitate the quantification of luciferase expression. The luciferase activity, as quantified via a luminescence reader (BioTek Synergy H1), revealed a dose-response relationship with varying concentrations of mating factor, demonstrating the threshold sensitivity and response kinetics of the bioparticles to the mating pheromone.

### Mating pheromone production quantification

Bioparticles were suspended in YPD medium to achieve an OD of 0.1, with 100 µl aliquots distributed into a 96-well plate. An initial dose of mating factor (IDT, custom peptide manufacturing) was added to the first well to reach a final concentration of 10 µM. A serial dilution was conducted across the subsequent wells by transferring 20 µl from one well to the next. The plate was incubated for 1 hour at 30°C in a shaking incubator (BioTek Synergy H1, 300 RPM). The plate was centrifuged at 1000 RCF for 1 minute in a swinging bucket centrifuge to remove the cellular fraction, and the resultant supernatants were employed for analysis in the Mating Pheromone Sensitivity Assay. The luciferase activity, as quantified via a luminescence reader (BioTek Synergy H1), delineated a dose-response relationship with varying concentrations of mating factor, demonstrating the mating pheromone production capacity of the bioparticles in response to the mating pheromone.

### Fold induction reporter experiments

Bioparticles were suspended in YPD medium to achieve an optical density (OD) of 0.1, with 100 µl aliquots dispensed into a 96-well plate. Four wells were treated with mating factor (IDT, custom peptide manufacturing) at a final concentration of 10 µM, alongside four other wells retained as noninduced controls, with both cohorts incubated at 30°C for a span of three hours. To establish blank values, four other wells were assessed immediately. Prior to assessing luminescence, OD600 readings were obtained for normalization. To separate cells from the medium, 100 µl of water was added to each well, followed by shaking the plate at 800 RPM for 3 minutes and centrifugation at 3000 RCF for 3 minutes. For readout, 11.4 ml of 0.1 M Tris (pH 8.6) was mixed with 300 µl of 1 µM furimazine (Promega, N1110) stock and 300 µl of 0.1% cycloheximide stock (Sigma-Aldrich, 01810). Subsequently, 15 µl of the supernatant from the centrifuged plate was mixed with 85 µl of readout buffer in a new 96-well plate, and luminescence was recorded over a span of 5 minutes using a luminescence reader (BioTek Synergy H1). The mean luminescence values were normalized against the initial OD600 readings to facilitate accurate interpretation of the data.

### Target cell fluorescent labeling

For adherent cell lines, cells were harvested utilizing StableCell solution (Sigma-Aldrich, T2601) following the manufacturer’s protocol, while suspended cell lines were employed directly. Cells were subsequently rinsed with fresh cultivation medium and treated with DNase I at 20 IU/ml (Sigma-Aldrich, DN25) for 5 minutes at 37°C. The ensuing cell suspension was filtered through a 20 µm filter (pluriSelect, 43-10020-60) and adjusted to a cellular concentration of 2.5×10^6^/ml. This suspension was then treated with ATTO425-maleimide dye (ATTO-TEC, AD 425-45) at a final concentration of 1 µM for 15 minutes in the dark. After treatment, the cells were rinsed twice with PBS + 0.1% gelatin (Sigma-Aldrich, G9391) and resuspended in the corresponding cultivation medium at a concentration of 1×10^6^/ml. The cell suspension was combined with a cryopreservation buffer (80% FBS, 20% DMSO; Sigma-Aldrich, F9665, D8418) in a 1:1 ratio, thoroughly mixed, and divided into aliquots. The resultant aliquots were placed in cryopreservation containers (Thermo Fisher Scientific, 5100-0001) and transferred to a −80°C freezer overnight incubation. After incubation, target cell aliquots were relocated to liquid nitrogen storage.

### Biocytometry workflow

Tubes containing BIOS Reagent (Sampling Human, Berkeley, CA, USA) were retrieved from −20°C storage and allowed to equilibrate to room temperature over a span of 3 minutes. Concurrently, target cell aliquots were retrieved from liquid nitrogen storage and thawed at 37°C. The target cell concentration was adjusted to the desired levels using Reaction Buffer (Sampling Human, Berkeley, CA, USA). Additional background cell lines were resuspended in Reaction Buffer and incorporated at specified concentrations where applicable. Reactions were prepared by adding 100 µl of the cell suspension to tubes with BIOS Reagent. Tubes were centrifuged at 50 RCF for 1 minute for a total of 10 cycles, rotating tubes by 180 degrees after each spin down. The resulting pellet was resuspended by agitation at 1400 RPM for 1 minute using a tube shaker (biosan, TS-100C). Subsequently, 75 µl of the reaction mixture was transferred to a tube with Hydrogel Medium (Sampling Human, Berkeley, CA, USA), calibrated to 30°C. Tubes were inverted to mix the contents, incubated at 37°C for 5 minutes, and inverted again. Reactions were then divided into aliquots and placed in a 96-well plate as needed. The plate was incubated at 4°C for 10 minutes, followed by a 4-hour incubation at 30°C. For readout, 11.4 ml of 0.1 M Tris (pH 8.6) was mixed with 300 µl of 1 µM furimazine (Promega, N1110) stock and 300 µl of 0.1% cycloheximide stock (Sigma-Aldrich, 01810). After incubation, a readout buffer was added, and luminescent readouts were acquired using a luminescence reader (BioTek Synergy H1). If applicable, the contents of the 96-well plate were analyzed via fluorescence microscopy (Olympus iX73 microscope, ×10 objective, CFP channel) to validate target cell counts in each well. The number of target cells in each well could then be validated against the corresponding luminescence measurement in each well.

### Data normalization

Luminescence readings from each well were averaged across five kinetic measurement points, generating a value denoted as *LUM_well_*. A baseline control value, abbreviated as *NC*, was established by averaging the luminescence output across eight wells specified as the negative control. Assuming the noise level to be 10% of the control value, the signal-to-noise ratio for each well, abbreviated as *SNR_well_*, was computed utilizing the formula:

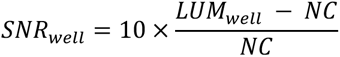

### Donor sample collection

Peripheral blood specimens were obtained from either healthy volunteers or patients undergoing stem cell mobilization regimens, adhering to protocols sanctioned by the Ethics Committee of the University Hospital in Pilsen (project BIODETEK, dated January 29, 2021). Vacutainer tubes with K_2_EDTA as the anticoagulant were used, with each sample averaging a volume of 7 ml. To mitigate endothelial cell contamination during venipuncture, the initial 2 ml of blood was discarded, a standard practice to ensure sample purity. After collection, samples were transported to the laboratory for processing and analysis within a 4-hour timeframe. In some instances, PBMCs were isolated using Ficoll gradient separation (described in subsequent sections). Bone marrow specimens were handled identically to peripheral blood samples, with no additional procedures undertaken.

### Ficoll gradient isolation

Peripheral blood mononuclear cells (PBMCs) were isolated from heparinized whole blood samples through a density gradient centrifugation technique. Initially, the whole blood was diluted in a 1:1 ratio with sterile phosphate-buffered saline (PBS) and carefully layered over a 6 ml volume of Ficoll-Paque (GE Healthcare, 17-1440-02) in 15 ml conical centrifuge tubes, ensuring minimal disturbance of the layers. The tubes were then centrifuged at 1000 RCF for 15 minutes at 15°C with a low brake and acceleration setting to maintain the integrity of the separated layers. Following centrifugation, the distinct mononuclear cell layer, formed at the interface of plasma and separation solution, was aspirated using a Pasteur pipette and transferred to a new 50 ml centrifuge tube containing 20 ml PBS. This cell suspension was then centrifuged at 350 RCF for 8 minutes, after which the supernatant was discarded, and the cell pellet was resuspended in a 3:2 PBS-albumin (Octapharma, 20% Albunorm) solution. Cell concentration was adjusted to 1×10^7^ cells/ml based on counts obtained from a Bürker chamber or hematological analyzer. For cryopreservation, a 0.5 ml aliquot of the cell suspension was mixed with an equal volume of chilled cryopreservation medium (DMSO + 20% albumin + PBS in a 1:2:2 ratio) and stored in cryovials, which were immediately placed in a pre-chilled cryopreservation container and transferred to a −80°C freezer for controlled rate freezing.

### Sample series preparation for peripheral blood and bone marrow

Peripheral blood or bone marrow samples were initially subjected to target cell enrichment, adjusting the target cell concentration to a specified level. Each sample contained the equivalent of 1×10^6^ leukocytes. After enrichment, a red blood cell (RBC) lysis solution (Biolegend, 420301) was employed to lyse red blood cells, adhering to the manufacturer’s protocol. After lysis, the samples underwent centrifugation at 350 RCF for a span of 3 minutes. The supernatant was carefully aspirated, leaving behind the cell pellet, which was then resuspended in DPBS supplemented with heparin (20 IU/ml; Sigma-Aldrich, H5515) to mitigate cellular aggregation. Each sample within a series was split into two, generating two identical sample sets designated for cellular analysis via biocytometry or flow cytometry.

***Preparation of sample series for PBMCs, primary cell cultures, and apoptotic primary cell cultures*** Peripheral blood mononuclear cells or primary cell culture samples were resuspended in DPBS supplemented with heparin (20 IU/ml, Sigma-Aldrich, H5515) to mitigate cellular aggregation. Samples were enriched with a target cell population, adjusting the target cell concentration to a specified level. Each sample within a series was split into two, generating two identical sample sets designated for cellular analysis via biocytometry or flow cytometry.

### Flow cytometry workflow

A 100 µl aliquot of cell suspension from each sample in the dilution series was transferred to a TruCount Absolute Counting tube (BD Biosciences, 340334). Following gentle vortexing, a specified antibody mix (all sourced from Biolegend, specific antibodies detailed in Supplementary Table 2) was introduced to the cell suspension. The antibody-cell mixture was then incubated for 15 minutes at 25°C in the dark to ensure optimal antibody binding. Cell analysis was conducted using a BD FACSCanto II flow cytometer (BD Biosciences). The flow cytometer was prepared for analysis through a washing step lasting for 30 minutes with distilled water and sheath fluid to eliminate any residual debris from previous runs. Prior to analysis, the instrument underwent daily calibration, encompassing optics, electronics, and fluidics tuning, alongside fluorescence compensation and detector voltage adjustment using BD CS&T beads (BD Biosciences). The data were collected, and compensation was carried out to correct for spectral overlap, employing Diva Flow Cytometry Analysis Software (BD Biosciences) to generate the final Flow Cytometry Standard (FCS) data file.

### Gate setting for flow cytometry analysis

The FCS files were exported as version 3.1 and subsequently converted to corresponding CSV files for analysis. Initially, utilizing a combination of FITC and PE channels, the dataset was filtered to exclude TruCount beads from the final sample analysis. Events showing nonspecific fluorescence (PB_A > 2000) were removed from the TruCount population to further refine the absolute event count calculations. Events exhibiting low FSC properties, characteristic of debris, were excluded, while high FSC and SSC events were retained to account for the target cell population’s tendency toward aggregation. The target cell population was then quantified using a high PB gate (PB_A > 3000) alone or in conjunction with a high APC gate (APC_A > 3000). Thresholds for positive event classification were determined using positive control samples, which contained a high ratio of target cell population within the respective sample matrix. Enumeration data for CD25+ or CD34+ populations were provided by our partner institution, who also conducted the corresponding analysis.

### Maintenance of primary cell cultures

Primary cell cultures of PBMCs were established following standard aseptic techniques, adhering to institutional biosafety and ethical guidelines. Post-isolation, PBMCs were counted using a Neubauer Improved hemocytometer, and viability was assessed via trypan blue exclusion (Sigma-Aldrich, 93595), ensuring a viability threshold of >95%. The cells were then seeded in a 12-well microplate at a density of 1×10^6^ cells/ml in RPMI 1640 medium (1 ml per well) supplemented with 10% heat-inactivated fetal bovine serum (Sigma-Aldrich, F9665) and 1% penicillin-streptomycin (Sigma-Aldrich, A5955). Cultures were maintained in a humidified incubator at 37°C with 5% CO_2_, with the medium refreshed every 48-72 hours.

### Apoptosis induction via heat shock

A 5 ml aliquot of the primary cell culture was carefully transferred into a sterile 15 ml Falcon tube, ensuring minimal cellular disturbance. The cell suspension was then subjected to heat stress by immersing the Falcon tube in a water bath set at 50°C for 30 minutes to induce apoptosis. Following heat exposure, the cell suspension was allowed to return to room temperature before being gently transferred back into the cultivation plate. The cultures were then placed in a 37°C incubator for a 6-hour recovery phase under standard cultivation conditions, a duration determined to allow for cellular response to the induced stress. After recovery, the cell suspension was used for downstream experimental applications.

### Activation induction via PHA

The PBMC primary cell culture was treated with phytohemagglutinin (PHA) at a final concentration of 2.5 µg/ml. The plate contents were homogenized using a standard plate shaker to ensure an even distribution of PHA across the culture. The plate was then placed in an incubation chamber maintained at 37°C with a 5% CO_2_ atmosphere for 48 hours, allowing optimal cellular activation. After the incubation period, the activated primary cell cultures were washed with fresh RPMI 1640 medium to remove residual PHA. The resulting cell suspension was used for further experimental evaluations.

### Annexin V staining

The cells were prepared for Annexin V staining by first undergoing a washing step with PBS + 0.1% BSA, followed by resuspension in Annexin V Binding Buffer (Biolegend, 422201) at a concentration of 1×10^6^ cells/ml. A 100 µl aliquot of the cell suspension was then transferred to a 1.7 ml microcentrifuge tube, to which 5 µl of FITC Annexin V was added. The cells were gently vortexed and incubated for 15 minutes at room temperature (25°C) in the dark. Following incubation, the cells were washed once in Annexin V Binding Buffer before analysis using fluorescence microscopy (Olympus iX73 microscope, ×10 objective, GFP channel).

### Preparation of HaCaT/Jurkat mixed cell cultures

The Jurkat and HaCaT cell cultures were retrieved postharvest and centrifuged at 250 RCF for 2 minutes. Subsequently, the supernatant was carefully discarded, and the cell pellet was resuspended in 1 ml of DMEM. To remove cell aggregates, the cells were sieved through a 20 µm filter (pluriSelect, 43-10020-60), and the cell density was adjusted to 4×10^5^ cells/ml. The coculture cultivation plate was prepared by dispensing 50 µl of the HaCaT cell suspension (4×10^5^ cells/ml) and 50 µl of the Jurkat cell suspension (4×10^5^ cells/ml) into the preselected wells. Gentle tapping of the plate was performed to facilitate a uniform cell distribution. For induction, the wells were treated with 2.5 µl of 100 mM DTT (Sigma-Aldrich, 43815), and the contents were mixed thoroughly using a pipette tip. The plate was incubated for 3 hours under standard cultivation conditions. After induction, the cell suspension was used for downstream experimental applications.

## Acknowledgements

We are grateful to SEKK s.r.o. for providing access to their external quality assessment datasets, including analyses of CD34+ cells.

## Data availability

The data that support the findings of this study are available from the corresponding author, D.G., upon reasonable request.

## Supplementary figures

**Supplementary Figure 1.**
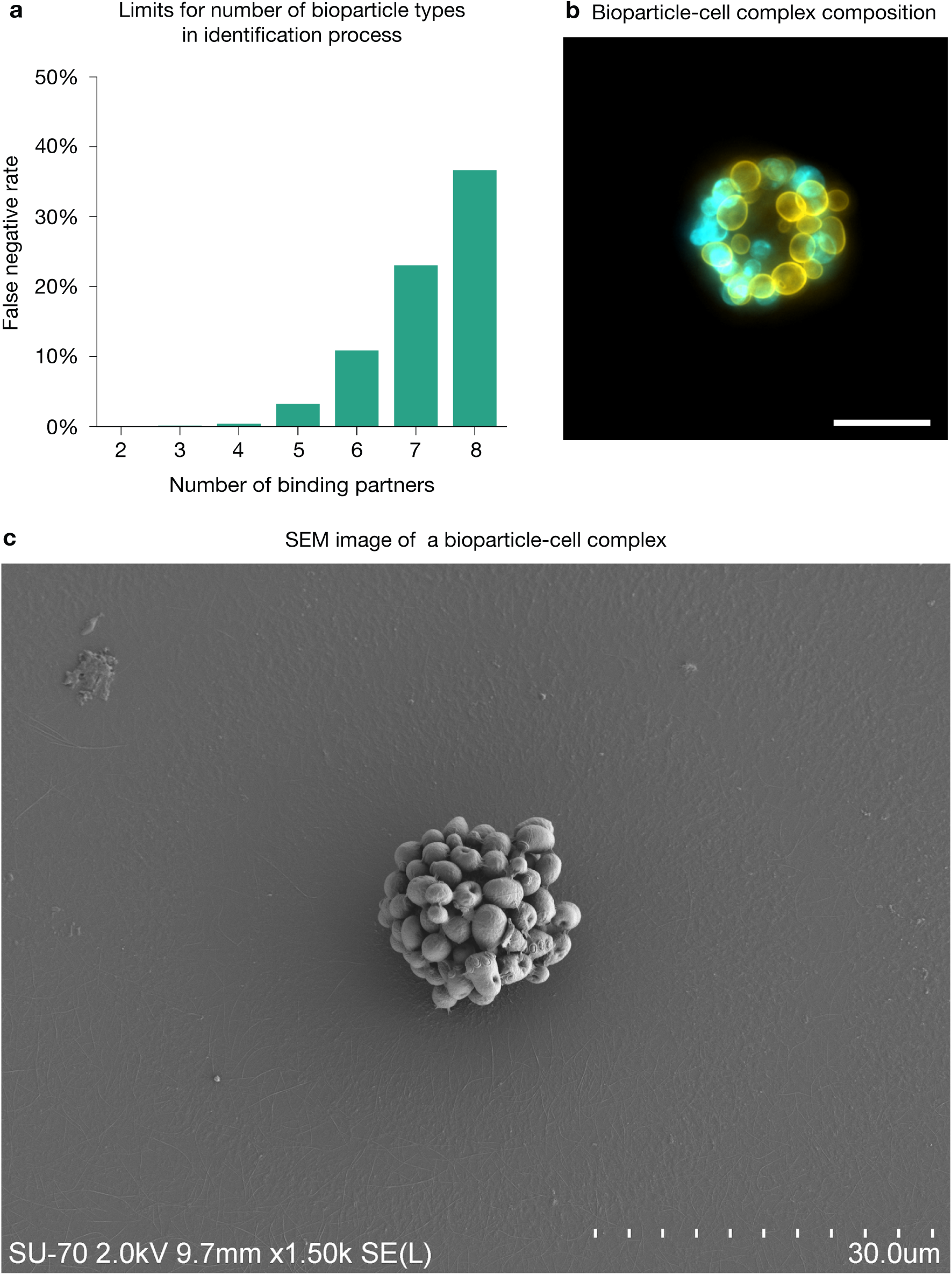
| Four surface markers may be processed during single identification event. (**a**) A statistical model demonstrates that up to four types of bioparticles, each with unique antigen specificity, can be used to determine a target profile with over 99% sensitivity. The assumptions of the model are as follows: a single target can support up to 60 bioparticles, corresponding to a target with an approximate radius of 10 µm (typical of a small mammalian cell). Each bioparticle is presumed to have an equal propensity for binding. It is also posited that at least six copies of a given bioparticle type are required to reconstruct a profile, a figure that has been experimentally validated (data not shown). (**b**) Equal representation of bioparticle types in bioparticle-cell complex was confirmed using the HaCaT cell line with the BIOS(EpCAM+EGFR) system: EpCAM-specific bioparticles stained blue and EGFR-specific bioparticles stained yellow. Binding conditions were optimized to ensure lower number of bound bioparticles, thus facilitating the differentiation of bioparticle types. The scale bar indicates 10 µm. (**c**) A scanning electron microscopy image facilitating the count of bioparticles on a target cell surface.

**Supplementary Figure 2.**
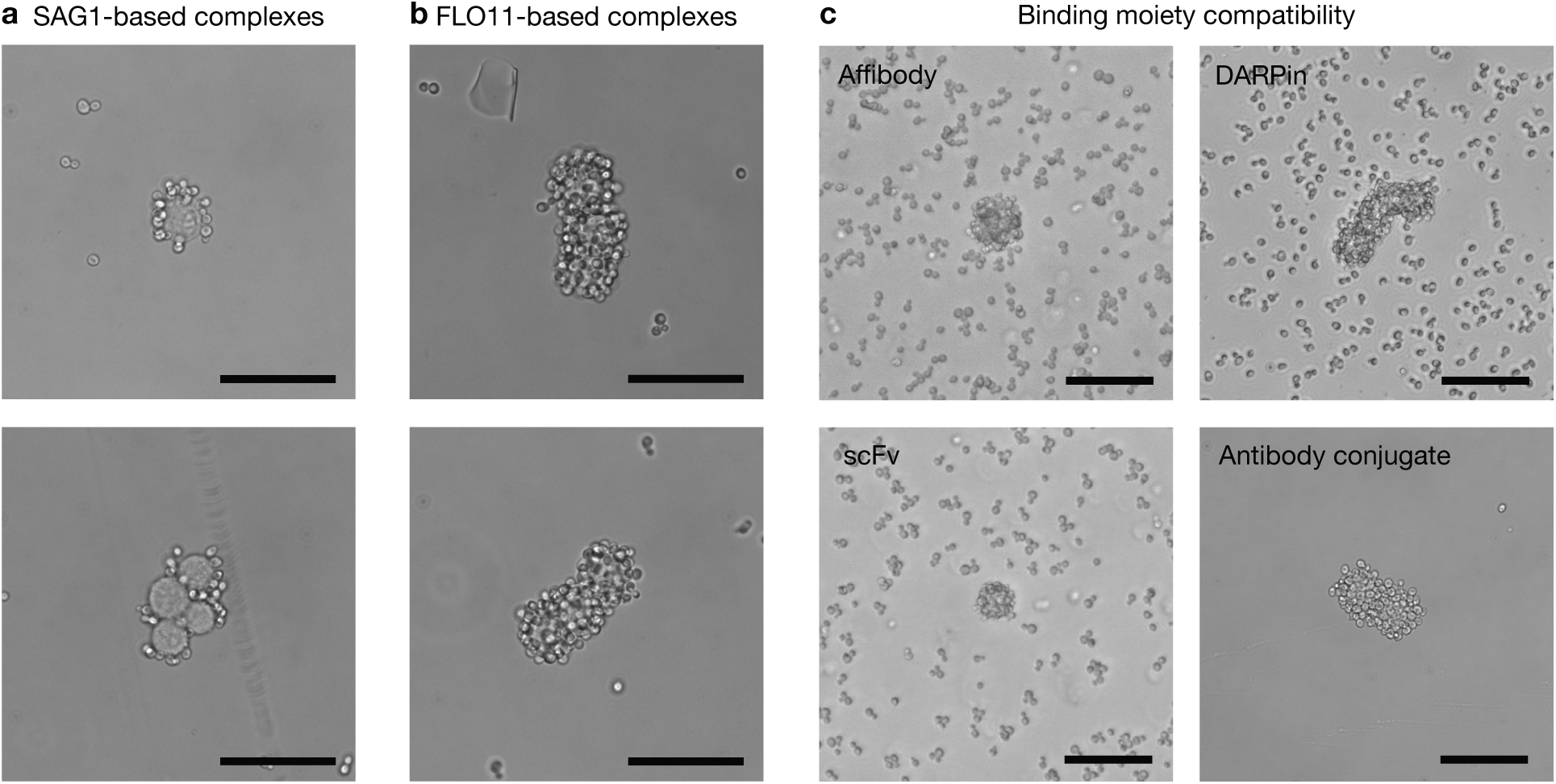
| Comparative analysis of bioparticle-cell complex formation using diverse anchoring systems. Evaluation of anchoring mechanisms on the effciency of bioparticle-cell complex formation. HaCaT cells (EpCAM+) were subjected to the standard biocytometry workflow using BIOS(EpCAM). Two different anchoring systems were employed by the bioparticle systems: the conventional SAG1 anchor (**a**) or the novel FLO11 anchor (**b**). From a randomly selected subset of 20 cell-bioparticle complexes per anchor type, comparative analysis revealed a substantial 652% enhancement in binding effciency when employing the FLO11 anchor system. (**c**) Diverse set of binders can support the formation of bioparticle-cell complexes. Microscopic visualization of complexes formed between BIOS utilizing anti-Her2 Affbody with HaCaT cells, anti-EGFR DARPin with HeLa cells, anti-CD45 scFv with HL-60 cells, and anti-CD71 antibody conjugate with Jurkat cells. The scale bar indicates 20 µm.

**Supplementary Figure 3.**
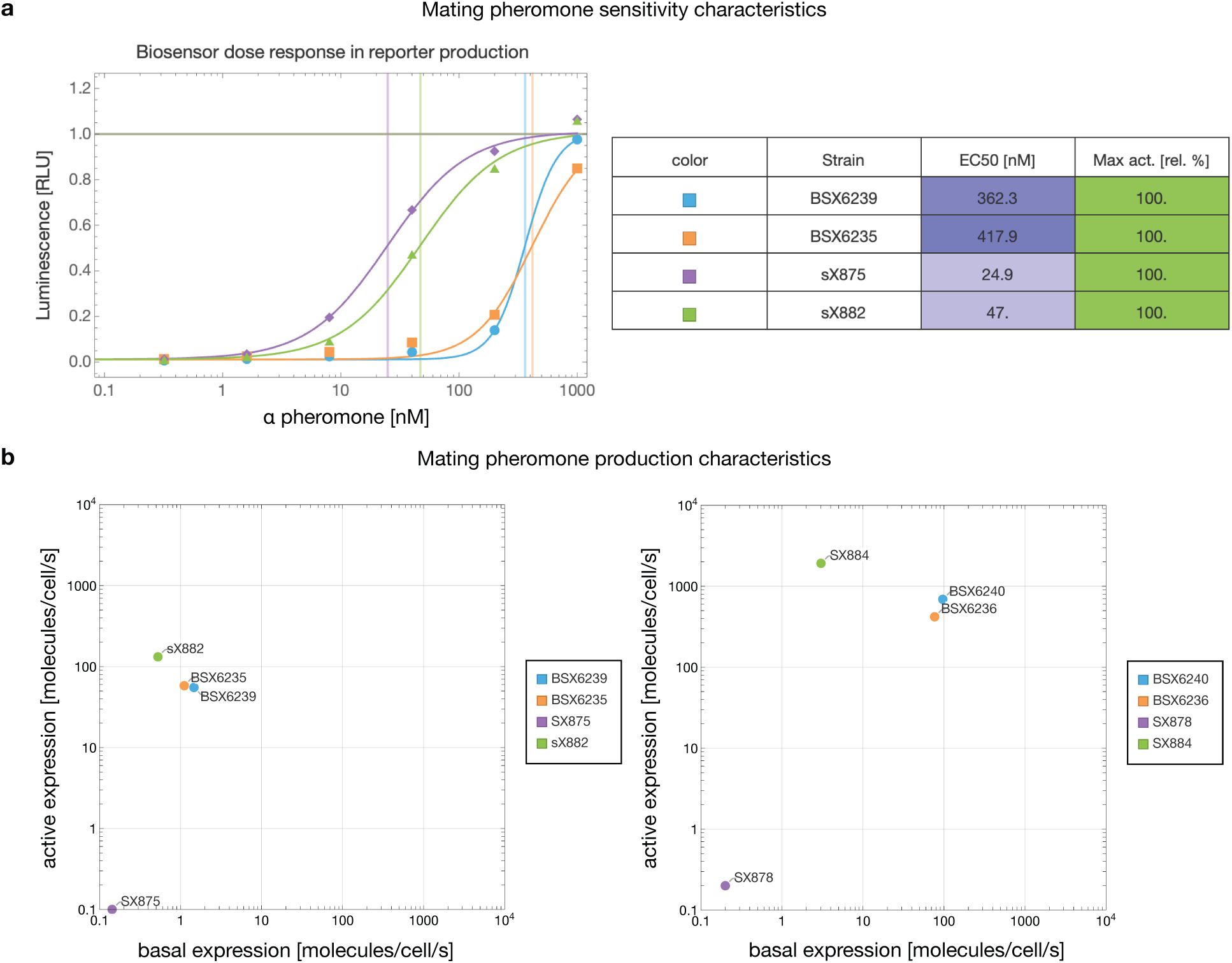
| Modulation of pheromone sensitivity and production capacity in engineered bioparticle strains. (**a**) Pheromone dose-response profiling of MATa-derived bioparticles was investigated across wild-type background with a reporter module (BSX6239), wild-type with reporter and binding modules (BSX6235), engineered background incorporating these modules (sX875), and strain with the full complement of reporter, binding, and signaling modules (sX882). Enhanced pheromone sensitivity is evidenced by the substantial reduction in EC50 in the engineered strains. (**b**) Quantitative analysis of pheromone basal and active production in bioparticles was investigated across wild-type backgrounds with a reporter module (BSX6239, BSX6240), wild-type with reporter and binding modules (BSX6235, BSX6236), engineered backgrounds incorporating these modules (sX875, sX878), and strains with the full complement of reporter, binding, and signaling modules (sX882, sX884). The results indicate minimized pheromone basal expression and enhanced production upon activation in the engineered strains.

**Supplementary Figure 4.**
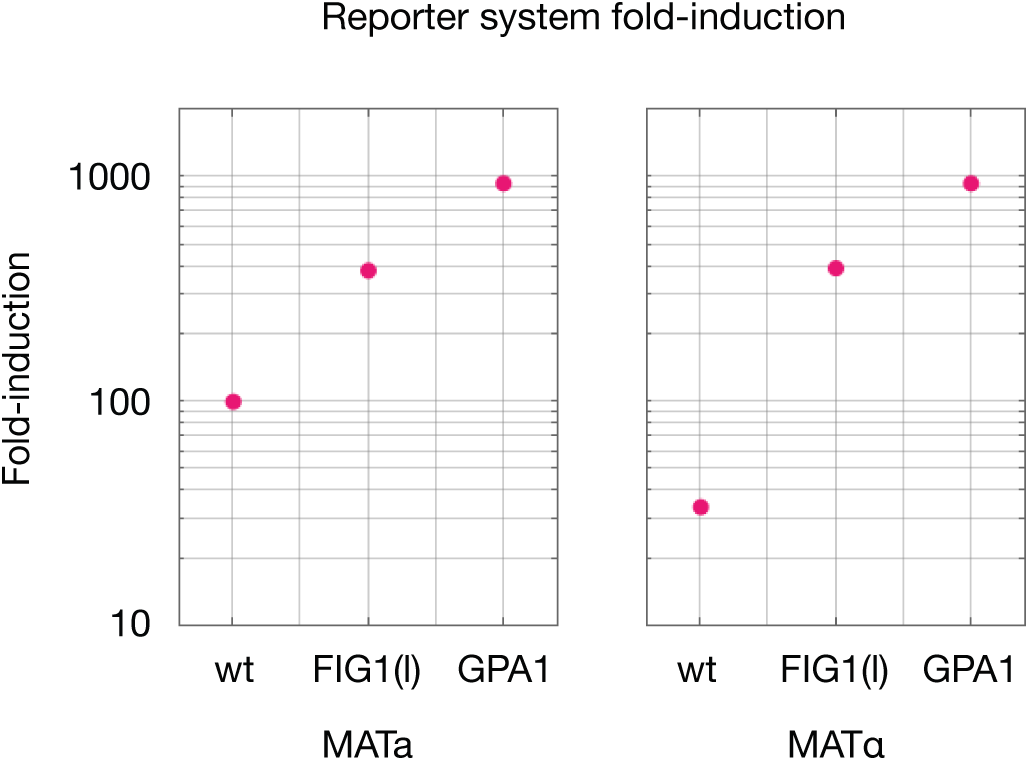
| Genetic modifications and their effects on reporter fold-induction. The reporter fold-induction was measured for three different bioparticle background strains to evaluate the effects of two genetic modifications: incorporation of *pFIG1(l)* and modification of the GPA1 subunit. For both modified strains, a significant increase in fold-induction was observed—a 10-fold increase for MATa-origin bioparticles and a 30-fold increase for MATα-origin bioparticles—with the *pFIG1(l)* implementation yielding the highest increase in reporter fold-induction for both mating types. The data presented are indicative of the ratio of reporter production measured at 180 minutes post-induction to the baseline reporter production without induction, across n = 2 biological replicates for each strain.

**Supplementary Figure 5.**
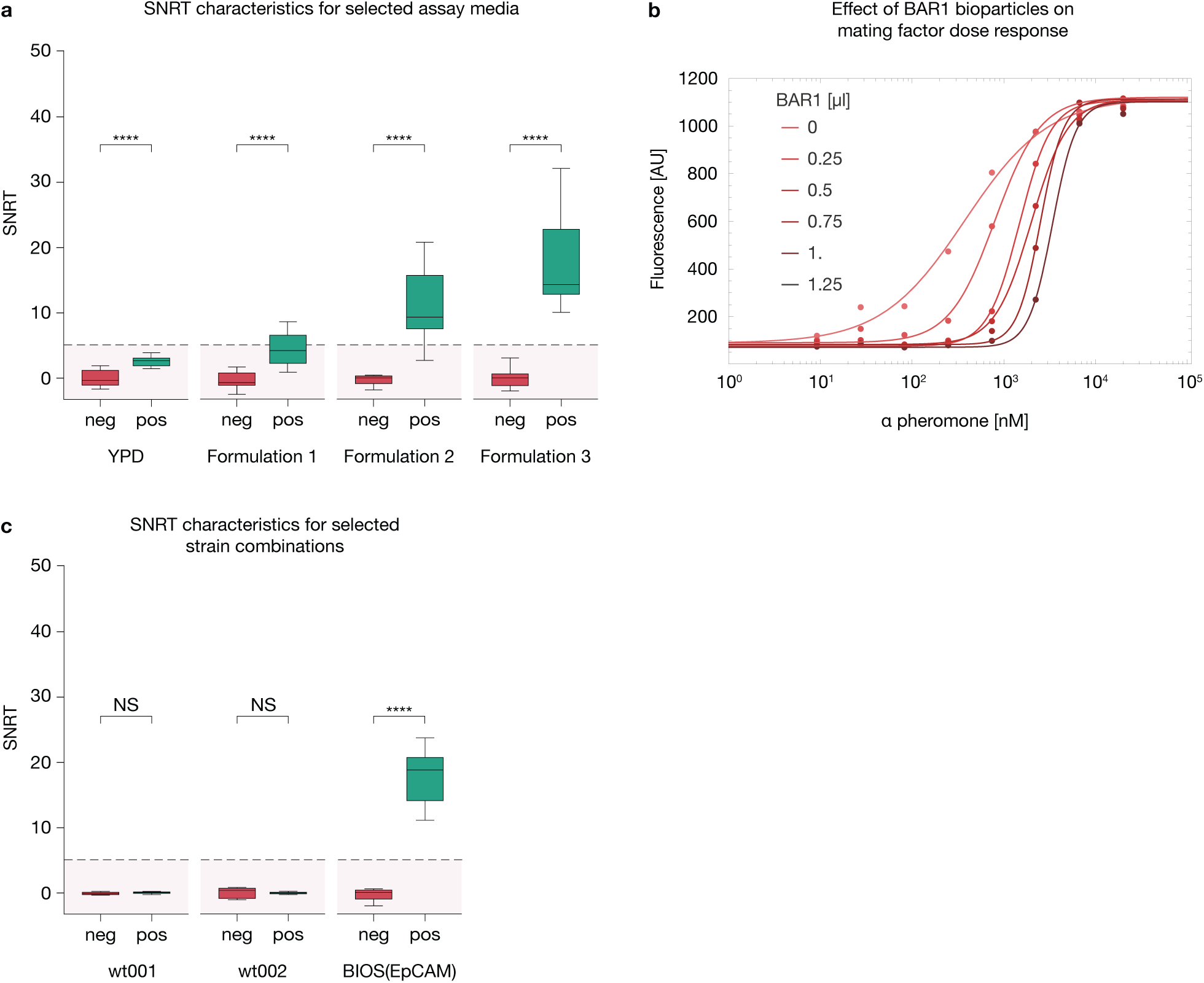
| Optimization and comparative analysis of assay media formulations for enhanced detection of target cells. (**a**) Assessment of assay media formulations on the detection effcacy of target HaCaT (EpCAM+) cells. Various media were tested, including YPD (the standard medium for yeast cultivation), Formulation 1 (comprising only suppressor biopolymers within a hydrogel matrix), Formulation 2 (containing only crowding biopolymers in a hydrogel matrix), and Formulation 3 (the hydrogel system employed throughout this publication). Sample tubes containing 0-30 target cells derived from HaCaT (EpCAM+) cell lines were analyzed using BIOS(EpCAM) per the standard protocol. The mean SNRT values obtained were 3.0 for YPD, 5.1 for Formulation 1, 12.8 for Formulation 2, and a notably higher 20.9 for Formulation 3. (**b**) The secreted protease in BAR1 bioparticles effectively degrades alpha-factor, raising the activation threshold of the biosensors. (**c**) Comparative analysis of assay performance using EpCAM-specific bioparticles: the wild-type S288C strain (wt001), a strain harboring a partial suite of chromosomal modifications (wt002), and the fully genetically modified BIOS(EpCAM) strain. The BIOS(EpCAM) strain outperformed with a mean SNRT value of 19.2, in contrast to 0.1 for wt001 and −0.2 for wt002. Error bars represent 1 s.d. ****, p < 0.0001; NS, nonsignificant. A line at SNR = 5 delineates the upper limit for negative samples.

**Supplementary Figure 6.**
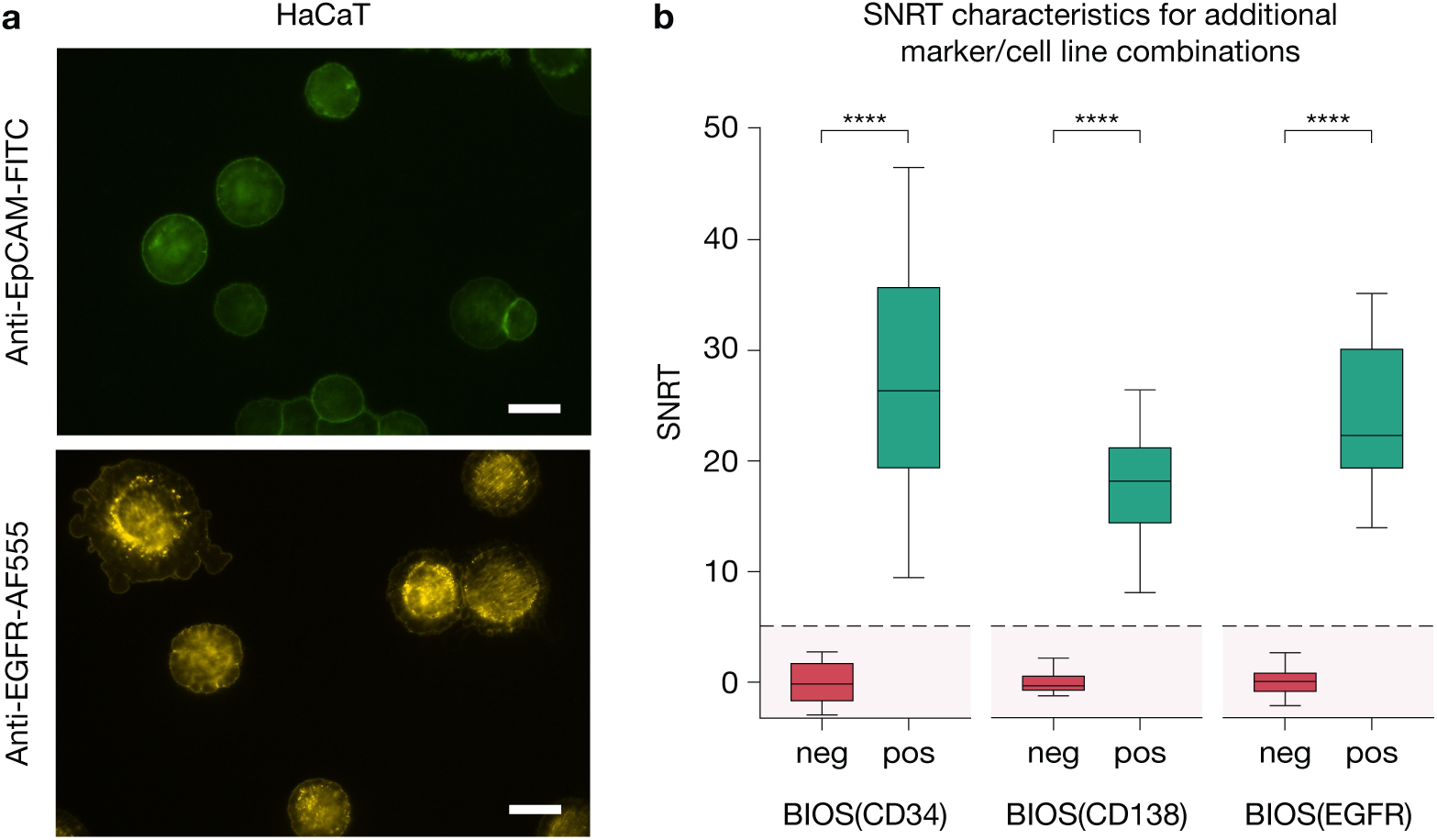
| Validation of cell surface marker detection and assay efficacy across cell lines. (**a**) EpCAM and EGFR expression in HaCaT cells was visualized using anti-EpCAM-FITC and anti-EGFR-AF555 antibodies, respectively. Microscopic examination confirmed a robust signal with correct membrane localization on HaCaT cells. The scale bar indicates 10 µm. (**b**) The effcacy of various BIOS systems was compared in assays with corresponding target cells: BIOS(CD34) with KG1a cells, BIOS(CD138) with RPMI8226 cells, and BIOS(EGFR) with HeLa cells. These combinations yielded consistent assay performance, with mean SNRT values of 26.7 for KG1a, 19.5 for RPMI8226, and 24.3 for HeLa cells. Error bars represent 1 s.d. ****, p < 0.0001; NS, nonsignificant. A line at SNR = 5 delineates the upper limit for negative samples.

**Supplementary Figure 7.**
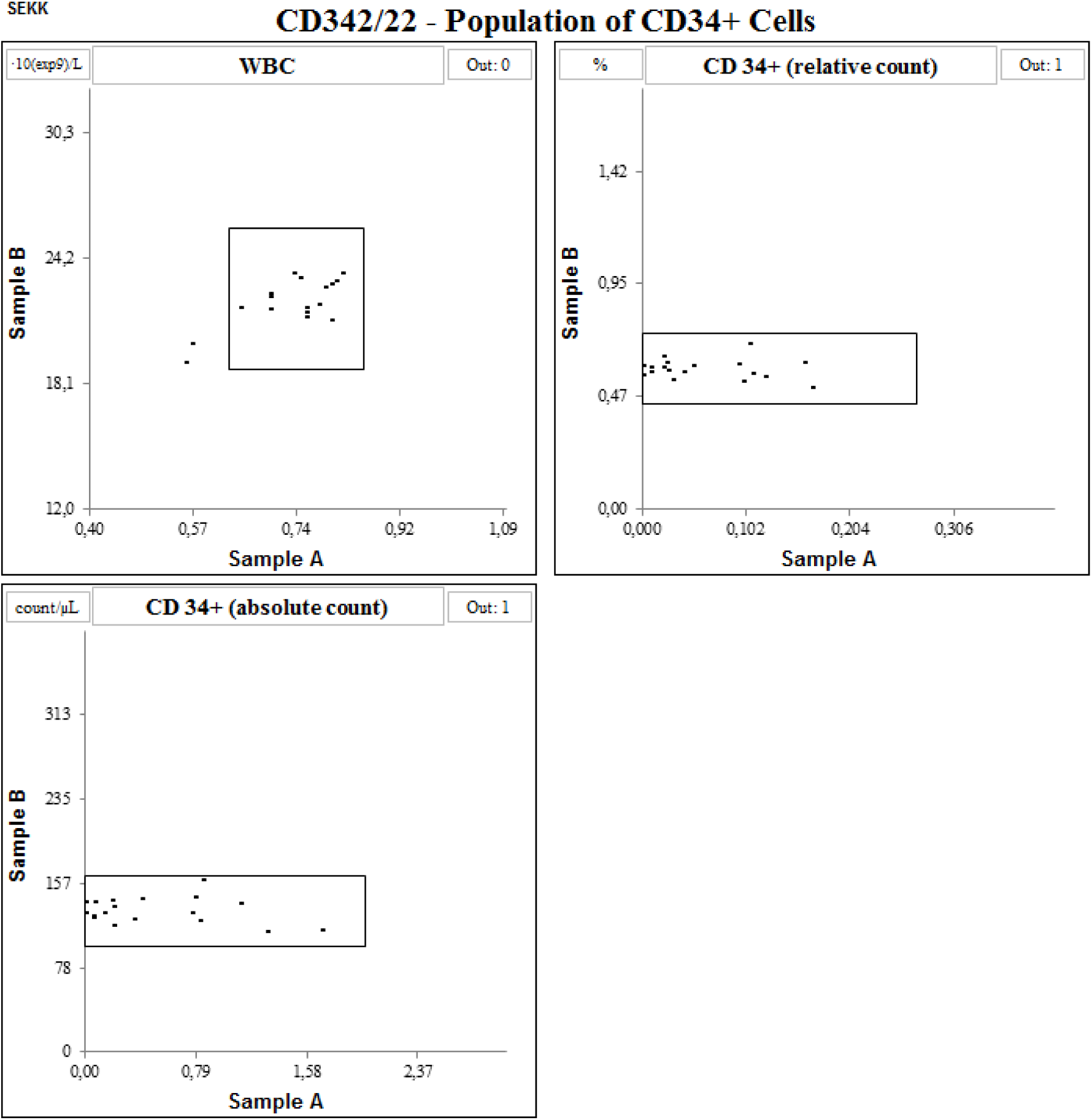
| Interfacility consistency of flow cytometry in CD34+ cell enumeration. Youden plots from a multicenter study display high interfacility concordance in quantifying cell populations larger than 0.5%. For cell counts approaching the detection limit of flow cytometry (∼ 0.1% cells), there is a notable increase in the coeffcient of variation, indicating variability and challenges in reproducibility. Reproduced with permission from SEKK s.r.o.

**Supplementary Table 1.**
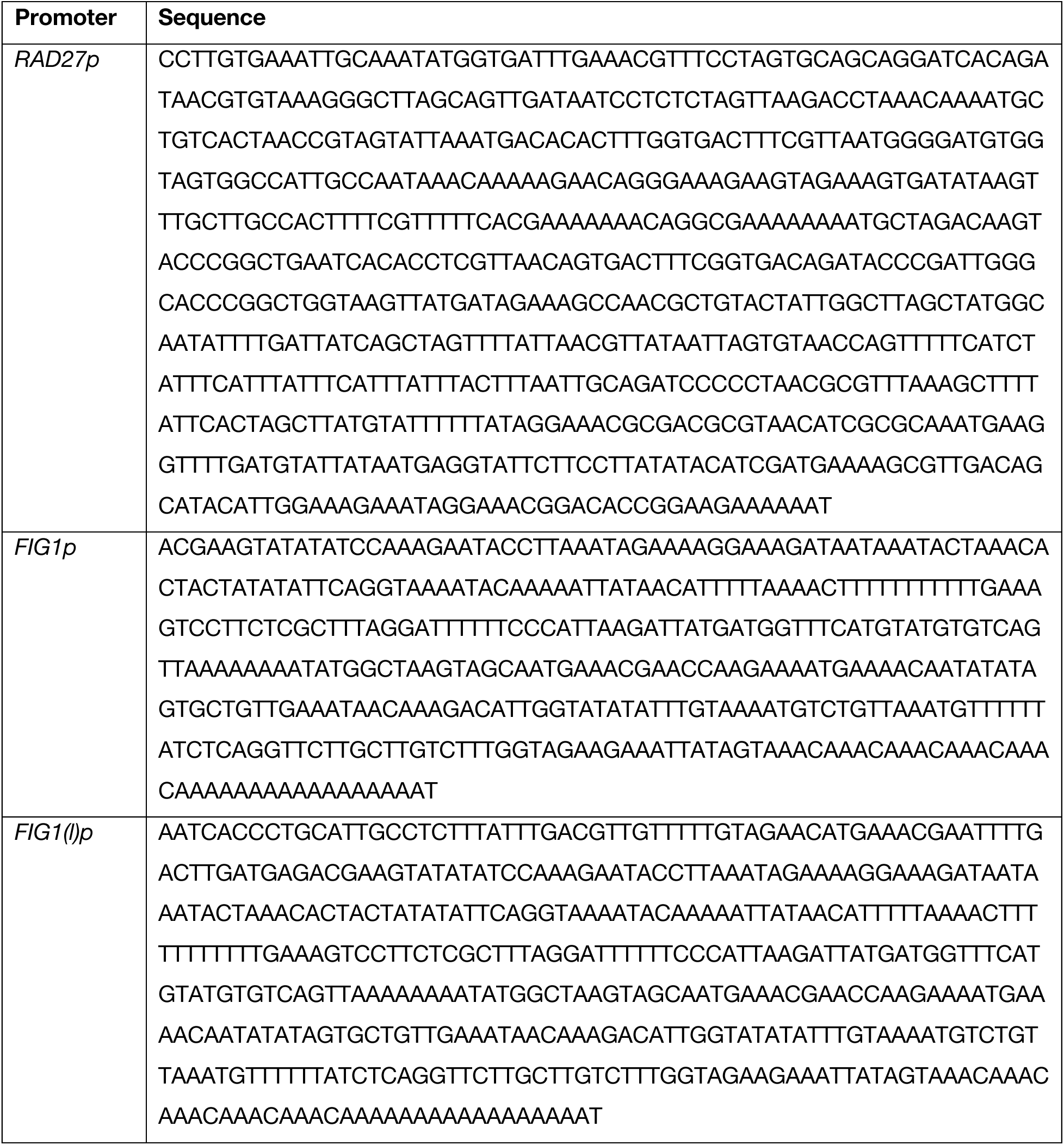
| Promoter sequences.

**Supplementary Table 2.** | Utilized antibodies.

| Target | Isotype | Conjugate | Clone | Product # |
| --- | --- | --- | --- | --- |
| EpCAM | Mouse IgG2b | Alexa Fluor 647 | 9C4 | 324212 |
| EGFR | Mouse IgG1 | Brilliant Violet 421 | AY13 | 352911 |
| CD25 | Mouse IgG1 | FITC | BC96 | 302604 |
| CD34 | Mouse IgG1 | PE | 581 | 343505 |

